# Compound models and Pearson residuals for single-cell RNA-seq data without UMIs

**DOI:** 10.1101/2023.08.02.551637

**Authors:** Jan Lause, Christoph Ziegenhain, Leonard Hartmanis, Philipp Berens, Dmitry Kobak

## Abstract

Recent work employed Pearson residuals from Poisson or negative binomial models to normalize UMI data. To extend this approach to non-UMI data, we model the additional amplification step with a compound distribution: we assume that sequenced RNA molecules follow a negative binomial distribution, and are then replicated following an amplification distribution. We show how this model leads to compound Pearson residuals, which yield meaningful gene selection and embeddings of Smart-seq2 datasets. Further, we suggest that amplification distributions across several sequencing protocols can be described by a broken power law. The resulting compound model captures previously unexplained overdispersion and zero-inflation patterns in non-UMI data.

## 1 Introduction

Single-cell RNA sequencing (scRNA-seq) data are affected by count noise and technical variability due to the total number of sequenced molecules varying from cell to cell. Removing this technical variation by normalization and variance stabilization is an important step in common analysis pipelines (Luecken and Theis, 2019; Heumos et al., 2023). The standard approach for this has been to use the log(1 + *x/s*) transformation, where *s* is a size factor of the cell. While the log-transform often performs well in practice (Ahlmann-Eltze and Huber, 2023), it has well-known theoretical limitations and can produce biased results (Lun, 2018).

Recently, a number of count modelling approaches like `sctransform` (Hafemeister and Satija, 2019), GLM-PCA (Townes et al., 2019), Sanity (Breda et al., 2021), and analytic Pearson residuals (Lause et al., 2021) have been suggested for pre-processing scRNA-seq data. These methods are based on explicit statistical models of the count generation process, rather than on heuristics such as the log-transform. One limitation of all of these methods is that they are tailored to data obtained using sequencing protocols based on unique molecular identifiers (UMIs), and are not appropriate for non-UMI technologies such as Smart-seq2 (Picelli et al., 2013). In this paper, we develop a count model and corresponding analytic Pearson residuals (Lause et al., 2021) for non-UMI sequencing data.

Single-cell sequencing protocols usually require an amplification step by polymerase chain reaction (PCR) to obtain enough starting material for sequencing (Figure 1). The process is imperfect, and different molecules will get amplified to a different extent. As a result, the number of sequenced molecules of a given gene (called *read count*) does not reflect the original number of RNA molecules in the cell.

**Figure 1:**
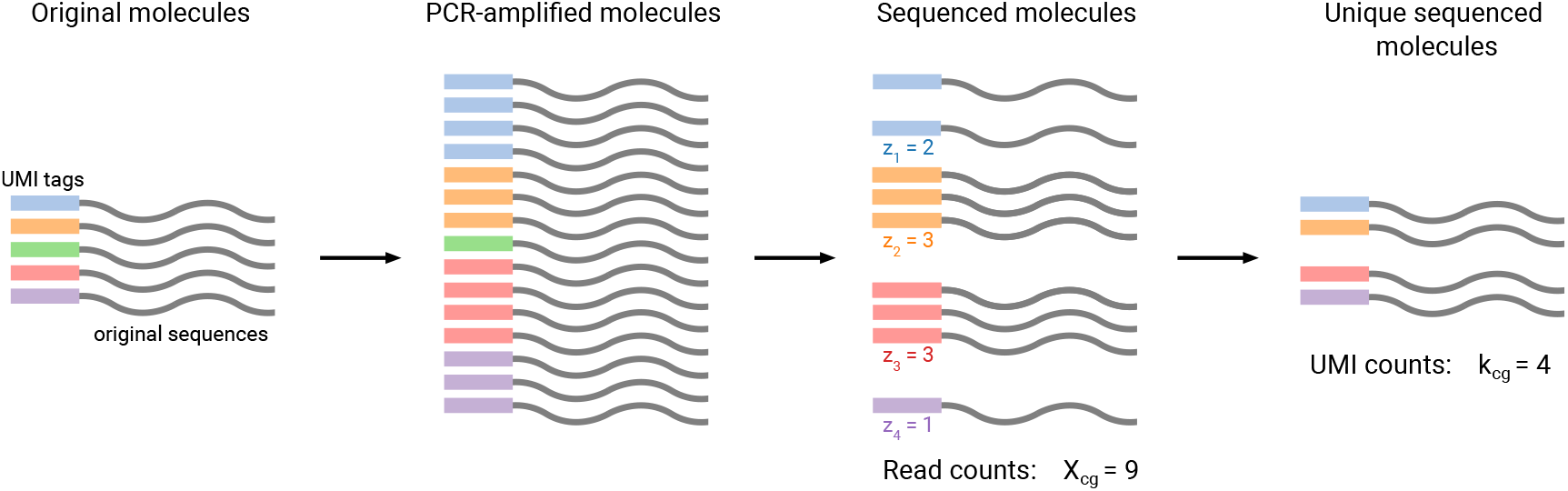
Important quantities in single-cell RNA sequencing. A cell *c* contains some number (here, 5) of RNA molecules of gene *g*. In UMI-based protocols, the original molecules are tagged by unique molecular identifiers (UMIs) before PCR amplification and sequencing. Both processes are imperfect, such that not all original molecules get amplified by the same factor, and not all amplified molecules get sequenced. In the end, each of the unique molecules may get sequenced one or several times, which is its *copy number* (*z*_*i*_). Here *z*_*i*_ values are 2, 3, 3, and 1, and one original molecule (green) is not detected at all. The sum of all copy numbers gives the observed *read count X*_*cg*_ (here, 9). UMIs allow to compute a *UMI count k*_*cg*_ by only counting unique sequenced molecules (here, 4). In non-UMI protocols, amplification and sequencing work the same, but the final de-duplication step is not possible, meaning that only *X*_*cg*_ is observable. Note that the shown numbers are not to scale: A typical cell might contain on the order of 10^5^ RNA molecules across all genes (Ziegenhain et al., 2022); yielding on the order of 10^10^ molecules after amplification; and producing on the order of 10^6^ sequenced reads.

In UMI protocols, a random DNA sequence called UMI is appended to each original reverse-transcribed RNA molecule prior to the amplification, and is then amplified and sequenced together with it (Islam et al., 2014). Because the UMI uniquely identifies each original molecule, one can later remove ampli-fication duplicates by counting each UMI only once (‘de-duplication’), giving rise to the *UMI counts* instead of *read counts* (Figure 1), effectively reducing amplification noise (Grün et al., 2014). Note that the UMI count does not necessarily equal the number of original RNA molecules present in the cell, as some molecules can get lost during sample preparation for the sequencing (‘library preparation’) or fail to get sequenced due to low capture rate (‘depth’) in the sequencing step (Figure 1).

While UMI protocols are popular in the scRNA-seq community, non-UMI technologies remain important. Indeed, UMIs only mark one end of the original molecules, so UMI counts are not available for the internal reads (most protocols involve a fragmentation step that cuts each molecule into small fragments prior to sequencing). Full-length sequencing methods, such as Smart-seq2 (Picelli et al., 2013), are typically more sensitive than UMI protocols (Ziegenhain et al., 2017; Ding et al., 2020), and are often used to detect rare cell types (Tasic et al., 2018; Yao et al., 2021) or splicing variants (Feng et al., 2021), or in low-throughput experiments such as Patch-seq (Lipovsek et al., 2021). In the resulting datasets, UMI counts are not available, and computational analysis has to be based on the read counts that still contain amplification-induced variability.

Only few normalization methods have been developed specifically to account for the amplification noise in read counts. The *Census* (Qiu et al., 2017) and *quasi-UMIs* (Townes and Irizarry, 2020) methods are two transformations that are designed to make the shape of the read count distribution approximately match the shape of the UMI count distribution. Afterwards, the transformed data still requires a UMI-like normalization. However, neither Census nor quasi-UMIs derive their transforms from a principled statistical model and rather rely on heuristics.

Here, we develop a new theoretically motivated method for normalization of non-UMI data that explicitly accounts for the amplification noise: *compound Pearson residuals*. We do so by extending the null model behind the analytic Pearson residuals (Lause et al., 2021) from UMI counts to read counts, based on an explicit statistical model for the amplification step. This yields a generative model for read counts that reproduces characteristic patterns of non-UMI data. We demonstrate that our compound Pearson residuals can efficiently normalize complex read count datasets.

## 2 Results

### 2.1 Analytic Pearson residuals for normalization of UMI data

In this section we briefly summarize the normalization approach based on Pearson residuals, originally developed by Hafemeister and Satija (2019) for UMI data. Pearson residuals compare the observed data to a null model that captures only technical variability due to count noise and variations in sequencing depth. The null model assumes perfect biological homogeneity, and so any deviation from it suggests biological variability.

Under the null model (Lause et al., 2021), a gene *g* takes up a certain constant fraction *p*_*g*_ of the total *n*_*c*_ RNA molecules sequenced in cell *c*, and the observed UMI counts *k*_*cg*_ follow a negative binomial (NB) distribution:

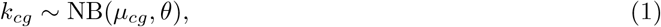

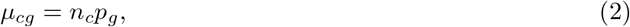

where *θ* is the inverse overdispersion parameter. Higher values of *θ* yield smaller variance, and for *θ* = ∞, the NB distribution reduces to the Poisson distribution. Note that *θ* in this formulation is shared between all genes; based on negative control UMI data, Lause et al. (2021) argued that *θ* can be set to *θ* = 100, which is close to Poisson. Sarkar and Stephens (2021) even suggest pure Poisson as measurement model.

Given an observed UMI count matrix, the maximum likelihood estimate of *µ*_*cg*_ is given by:

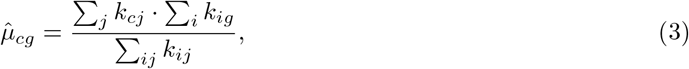

which is exact in the Poisson case and holds only approximately in the NB case (Lause et al., 2021). This yields the analytic formula for *UMI Pearson residuals* (difference between observed UMI count values and model prediction, divided by the model standard deviation):

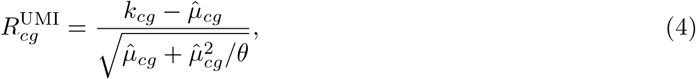

where 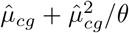 is the variance of the NB distribution with mean 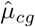 and overdispersion parameter *θ*. The variance of Pearson residuals does not depend on *p*_*g*_, and in a homogeneous dataset is close to 1 for all genes. This ensures variance stabilization across all levels of gene expression.

This algorithm is similar to the one implemented in sctransform (Hafemeister and Satija, 2019; Choudhary and Satija, 2021) and is equivalent to a rank-one GLM-PCA (Townes et al., 2019), but it is simpler and faster to compute than either of these methods. When followed by singular value decomposition (SVD), the Poisson (*θ* = ∞) version of UMI Pearson residuals is also known as correspondence analysis (Hsu and Culhane, 2023).

### 2.2 Compound Pearson residuals for non-UMI read count data

To apply Pearson residuals to scRNA-seq data without UMIs, we need to change the null model, because read counts do not follow the NB distribution (Svensson, 2020; Cao et al., 2021). As in the UMI case, we assume that the number of unique sequenced RNA molecules *k*_*cg*_ follows a Poisson or a NB distribution. However, during the amplification step, each of these *k*_*cg*_ unique molecules could have been duplicated multiple times before sequencing (Figure 1). For the *i*-th unique molecule, we call the number of its sequenced duplicates its *copy number z*_*i*_. We assume that copy numbers follow some distribution Z, which we call *amplification distribution*. Our assumption is that the amplification distribution is the same for all genes and all cells, and only depends on the details of the PCR amplification and the sequencing protocol (see the note below about the variable gene length).

The read count *X*_*cg*_ of a given gene *g* in cell *c* is thus modeled as the sum of *k*_*cg*_ independent and identically distributed (i.i.d.) positive integer copy numbers drawn from Z:

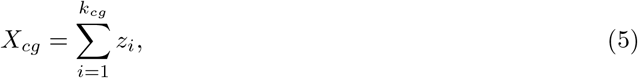

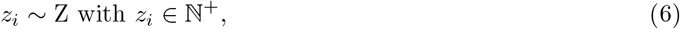

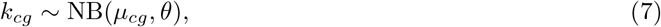

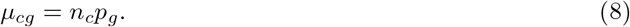

For example, *k*_*cg*_ = 4 means that four unique RNA molecules of gene *g* were sequenced in a cell *c*; if their copy numbers were 2, 3, 3, and 1, this would yield the read count value *X*_*cg*_ = 2 + 3 + 3 + 1 = 9 (c.f. Figure 1). The resulting distribution of *X*_*cg*_ can be called *compound NB distribution* (see Methods).

In the above formulation, our model does not explicitly account for gene length. In most sequencing protocols, longer transcripts are cut into more fragments before amplification. This will result in more unique sequenced fragments *k*_*cg*_ for longer molecules (Phipson et al., 2017). In our model, this increase amounts to a constant length factor *l*_*g*_ per gene, which can be absorbed by our per-gene expression fraction *p*_*g*_. The variance of Pearson residuals does not depend on *p*_*g*_ (see above), so for simplicity, we do not explicitly put gene lengths into the model.

To obtain Pearson residuals for this null model, we need to obtain expressions for its mean and variance. The mean of a compound NB distribution is equal to the product of the NB mean *µ*_*cg*_ and the mean of the amplification distribution Z:

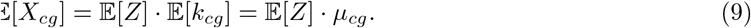

We can use the observed read count matrix *X* to estimate the means of the maximum likely null compound model as follows:

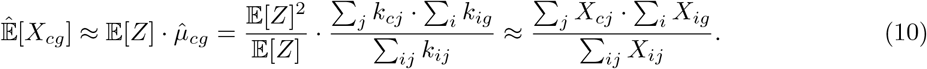

Note that this expression has the same form as Equation 3: the outer product of the row and the column sums of the count matrix, normalized by its total sum.

The mean-variance relationship of the compound NB distribution takes the form (see Methods):

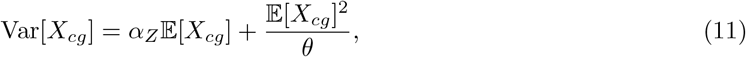

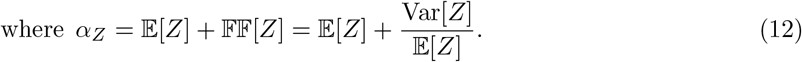

This expression is similar to the mean-variance relationship of the NB distribution but contains a scaling parameter *α*_*Z*_ equal to the sum of the mean and the Fano factor (denoted FF[*Z*]) of Z. Note that in the compound Poisson case (*θ* = ∞), the compound variance is proportional to the compound mean.

Using these equations, we can compute the Pearson residuals of the compound NB null model, which we call the *compound Pearson residuals*:

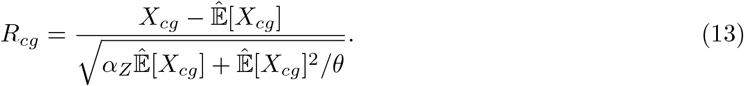

Following the UMI case and the arguments in Lause et al. (2021), we set the overdispersion parameter to *θ* = 100. The scalar *α*_*Z*_ is a function of the mean and variance of the amplification distribution and remains as the only free parameter of the model. Following Hafemeister and Satija (2019) and Lause et al. (2021), we clip the residuals to 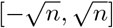, where *n* is the number of cells in the dataset.

This formalism naturally generalizes the UMI Pearson residuals. Indeed, in the UMI case, each sequenced molecule is counted only once, thanks to the UMIs, meaning that the Z distribution is a delta peak *δ*(1) with 𝔼 [*Z*] = 1 and Var[*Z*] = 0, and hence *α*_*Z*_ = 1. In this case, Equation 13 reduces to Equation 4.

Conveniently, compound Pearson residuals are equivalent to UMI Pearson residuals of the read count matrix scaled by 1*/α*_*Z*_:

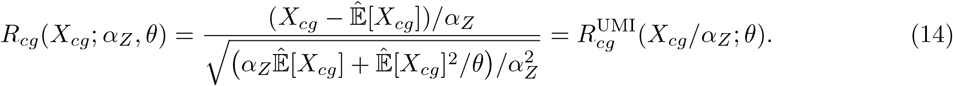

Importantly, the necessary scaling factor is not equal to 𝔼 [*Z*], as could be naïvely expected, but rather to *α*_*Z*_ = 𝔼 [*Z*] + Var[*Z*]*/ 𝔼* [*Z*].

Compound Pearson residuals have the same computational complexity as UMI Pearson residuals, and can be computed in seconds even for large datasets with *>*10 000 cells. The matrix of Pearson residuals is dense and will thus require more memory than the sparse matrix of raw counts. For very large datasets it may be prohibitive to hold the full matrix in memory, but memory demand can be reduced by advanced implementations (Irizarry, 2021) or by subsetting to highly variable genes (Lause et al., 2021), allowing to process datasets with millions of cells.

### 2.3 Compound model can fit homogeneous read count data

The compound model introduced above is designed to capture only technical, but not biological variance in non-UMI read count data. Therefore, it should provide a good fit to data that contain little biological variation. To test this, we took scRNA-seq data from adult mouse neocortex sequenced with Smart-seq2 (Tasic et al., 2018), and focused on a subset of cells corresponding to one specific cell type, assuming that there is little biological variability within a cell type. We chose the *L6 IT VISp Penk Col27a1* type (as annotated by the authors of the original study), containing 1 049 excitatory neurons (Figure 2).

**Figure 2:**
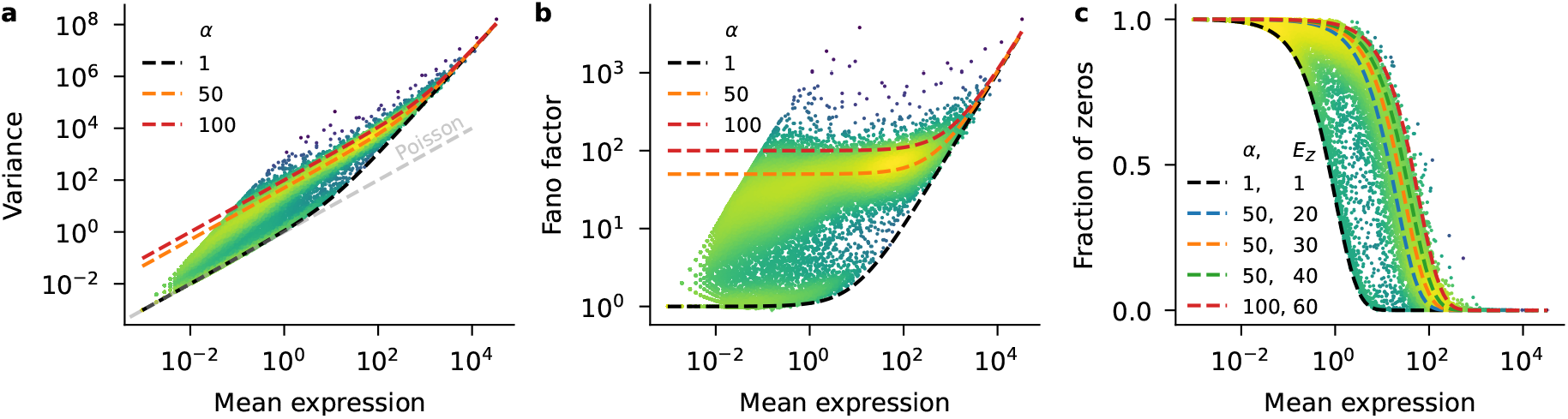
The compound NB model captures statistics of homogeneous read count data. The data are a homogeneous subset of a mouse visual cortex dataset (Tasic et al., 2018) sequenced with Smart-seq2 (*L6 IT VISp Penk Col27a1* cluster; 1 049 cells, 33 914 genes). Each dot represents a gene. Brighter colors indicate higher density of points. Dashed lines show the behavior of the compound negative binomial model (*θ* = 10). **a:** Meanvariance relationship. Gray line illustrates the Poisson case where mean equals variance. **b:** Relationship between mean expression and Fano factor (variance/mean). **c:** Relationship between mean expression and fraction of zero counts.

The mean-variance relationship across genes (Figure 2a) showed that most genes exhibited overdispersion compared to the Poisson model (gray line, *α*_*Z*_ = 1 and *θ* = ∞). Most genes also showed more variance than expected from a NB model without amplification (black line, *α*_*Z*_ = 1 and *θ* = 10). In contrast, compound models accounting for amplification with *α*_*Z*_ ∈ [10, 100] (colored lines) were able to approximate the mean-variance relationship for the majority of the genes. Note that we used *θ* = 10 for illustrations in Figure 2 because this value fit the within-cell-type data better than *θ* = 100, in agreement with the idea that even biologically homogeneous data can show some additional variability on top of the purely technical variability (Lause et al., 2021). Cell-to-cell variation in sequencing depth also contributed to this increase in overdispersion (Supplementary Figure S1).

The relationship between the mean and the Fano factor across genes (Figure 2b) allowed us to further constrain the amplification parameter *α*_*Z*_. Indeed, for genes with low average expression, the Fano factor of the read counts is approximately equal to *α*_*Z*_ (Equation 11 and Equation 29 in the Methods). The Fano factors of most genes were bounded by models with *α*_*Z*_ = 1 from below (black), and by models with *α*_*Z*_ = 100 from above (red). The bulk of the genes followed a model with *α*_*Z*_ = 50 (orange).

While knowing *α*_*Z*_ is sufficient to compute Pearson residuals, we can obtain separate estimates of 𝔼 [*Z*] and Var[*Z*] by studying the relationship between the average expression and the fraction of zeros. This relationship only depends on 𝔼 [*Z*] (see Methods, Equation 30), allowing to estimate this term directly. We observed that in the Smart-Seq2 data, the fraction of observed zeros decreased with increasing mean expression (Figure 2c). There were more observed zeros than expected from a NB model with *α*_*Z*_ = 1 (black), hinting at why read count data have in the past often been modeled using a zero-inflated negative binomial (ZINB) distribution (e.g. Lopez et al., 2018; Chen et al., 2018). Our compound NB model with 𝔼 [*Z*] ≈ 30 (orange) provided a good qualitative fit to the observed data, without any explicit zero-inflation terms. From here we can compute 𝔽 𝔽 [*Z*] = *α*_*Z*_ − 𝔼 [*Z*] ≈ 20 and hence Var[*Z*] ≈ 600.

The compound NB model with *α*_*Z*_ = 50 and 𝔼 [*Z*] = 30 described the majority of genes well. However, some genes were instead following the model without any amplification (black line in Figure 2), as if their transcripts were not amplified by the PCR. To understand this pattern, we obtained gene type annotations from `mygene.info`. This revealed that protein-coding genes generally followed our best-fitting compound model (Supplementary Figure S2a–c), while most of the seemingly non-amplified genes were pseudogenes (Supplementary Figure S2d–f). This observation was not limited to the Smart-seq2 data, but also occurred for all sequencing protocols studied in Ziegenhain et al. (2017) (Supplementary Figure S3). Exonic transcript lengths from the `mygene.info` database were shorter for the non-amplified genes (Supplementary Figure S4).

In summary, we showed that our compound NB model fits a biologically homogeneous example dataset. In particular, our model matched the main statistical properties of protein coding genes (mean, variance, and fraction of zeros).

### 2.4 Compound Pearson residuals for normalization of heterogeneous read count data

Next, we computed compound Pearson residuals with *α*_*Z*_ = 50 (Equation 13) to preprocess the entire dataset from Tasic et al. (2018), which is highly heterogeneous and includes both neural and non-neural cells from two areas of the mouse neocortex.

For highly variable gene (HVG) selection, we used the variance of compound Pearson residuals for each gene (Figure 3a). Most genes had residual variance close to 1, indicating that they followed the null model. The interpretation is that those genes did not show biological variability and were not differentially expressed between cell types. In contrast, genes with residual variance ≫ 1 had more variability than predicted by the null model, implying nontrivial biological variability. For downstream analysis, we selected the 3 000 HVGs with highest residual variances. Among those were well-known marker genes of specific cortical cell types (Figure 3a, red dots). In particular, the four genes with highest residual variance (star symbols in Figure 3a) marked different groups of inhibitory neurons (*Npy, Vip, Sst*) and astrocytes (*Apoe*). As before, we confirmed that genes with very low residual variances ≪ 1 were mostly pseudogenes (Supplementary Figure S5).

**Figure 3:**
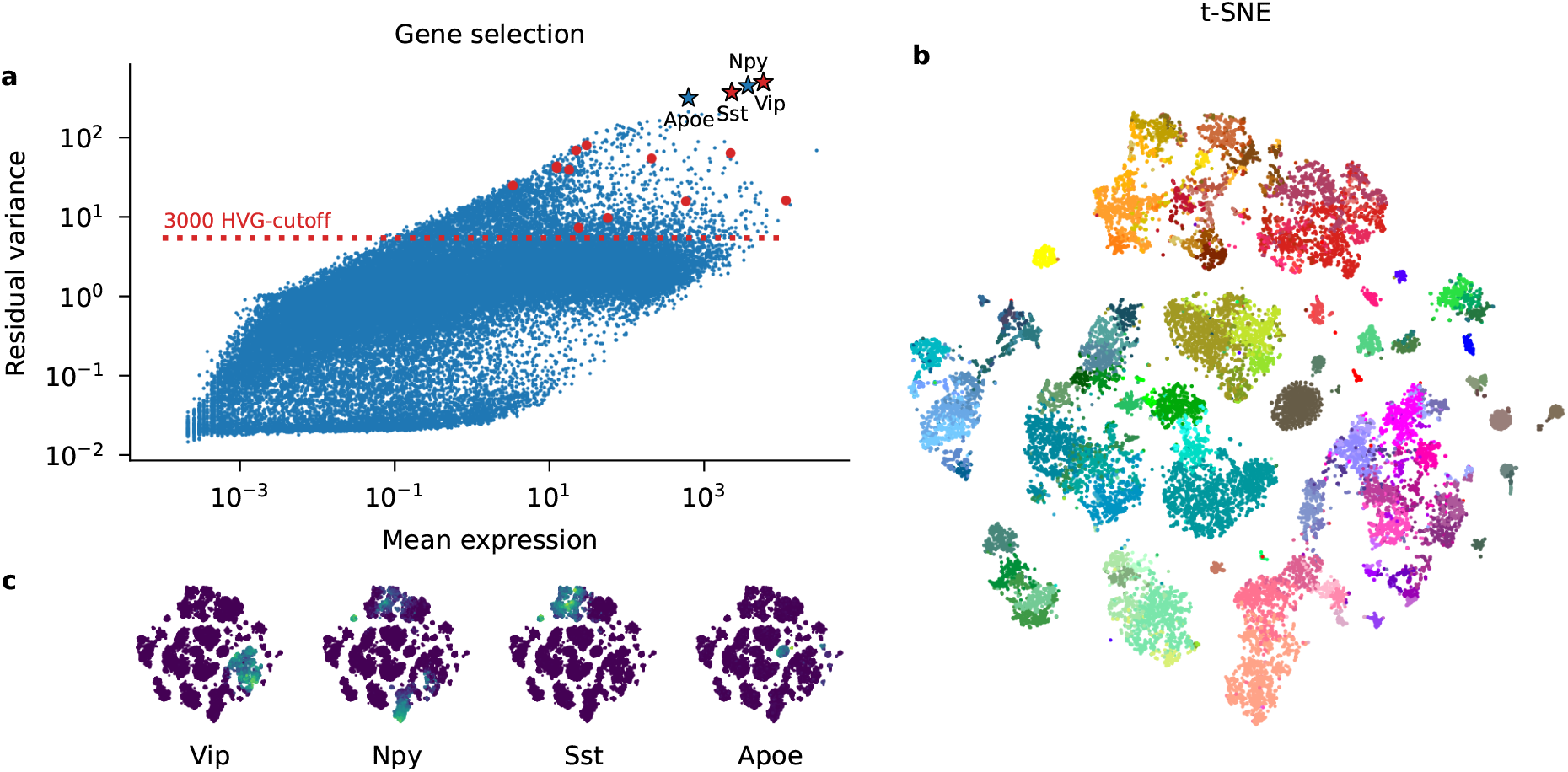
Compound Pearson residuals work well for preprocessing a heterogeneous Smart-seq2 dataset. Here, we used the raw counts of a mouse visual cortex dataset sequenced with Smart-seq2 (Tasic et al., 2018) (23 822 cells, 38 510 genes). We used a compound NB model with amplification parameter *α* = 50 and overdispersion parameter *θ* = 100. **a:** Highly variable gene (HVG) selection by largest residual variance. Each dot is a gene; genes above the red line were included in the selection of 3 000 HVGs. Stars indicate the top four HVGs shown in panel (c). Red dots and stars correspond to the following well-known marker genes taken from Tasic et al. (2016), from left to right: *Itgam* (microglia), *Bgn* (smooth muscle cells), *Pdgfra* (oligodendrocyte precursors), *Aqp4* (astrocytes), *Flt1* (endothelial cells), *Foxp2* (layer 6 excitatory neurons), *Mog* (oligodendrocytes), *Rorb* (layer 4 excitatory neurons), *Pvalb* (subset of inhibitory neurons), *Slc17a7* (excitatory neurons), *Gad1* (inhibitory neurons), *Sst* (subset of inhibitory neurons), *Vip* (subset of inhibitory neurons), *Snap25* (neurons). **b:** t-SNE embedding on compound Pearson residuals following HVG selection (to 3 000 HVGs) and PCA (down to 1 000 PCs). Each dot is a cell, colored by the original cluster assignments from Tasic et al. (2018). Warm colors: inhibitory neurons. Cold colors: excitatory neurons. Brown and gray colors: non-neural cells. **c:** t-SNE embeddings as in panel (b), colored by expression strength of the four most variable genes according to compound Pearson residual variance. For expression, we show square-root-transformed, depthnormalized counts.

Next, we used PCA and t-SNE to visualize the single-cell composition of the mouse cortex using compound Pearson residuals of the selected HVGs. The resulting embedding showed rich structure that corresponded well to the cell type annotations originally determined by Tasic et al. (2018) (Figure 3b): individual cell types formed mostly clearly delineated clusters, while related cell types (having similar colors) mostly stayed close to each other. The expression of most variable genes according to the residual variance was typically localized in one part of the embedding space (Figure 3c).

Calculating compound Pearson residuals requires to set the amplification parameter *α*_*Z*_. To investigate the influence of this parameter, we computed compound Pearson residuals for a range of *α*_*Z*_ values covering three orders of magnitude (Supplementary Figure S6). We found that for *α* ≫ 1, the exact value did not lead to large differences in the HVG selection or t-SNE representation. In contrast, when we used UMI Pearson residuals of the null model without amplification (*α*_*Z*_ = 1), the HVG selection failed to include some of the most important marker genes (Supplementary Figure S6a) and the embedding quality visibly degraded (Supplementary Figure S6b). This shows that it is not appropriate to apply the original formulation of UMI Pearson residuals (Lause et al., 2021) to non-UMI data, and that it is important to explicitly account for the PCR-induced variance. Reassuringly, the exact value of the *α*_*Z*_ parameter did not have a large influence on the downstream performance.

We compared our approach to existing methods for read count normalization: qUMI (Townes et al., 2019) and Census (Qiu et al., 2017). Both use heuristics to estimate UMI counts from read count data, and one can then apply standard UMI methods for further processing. We found that both methods, when combined with UMI Pearson residuals, gave results that were similar to our compound Pearson residuals (Supplementary Figure S7). This is unsurprising, as Census amounts to dividing read counts by a cell-specific constant, and compound Pearson residuals are equivalent to UMI Pearson residuals after appropriate scaling of the data matrix (Equation 14). The qUMI transformation is non-linear but gave similar results for our data. Importantly, both Census and qUMI rely on heuristics (see Discussion), while our approach is based on an explicit statistical model.

We also compared this approach to the default preprocessing implemented in the `Scanpy` library (Wolf et al., 2018) based on depth normalization, `log1p()` transform, and `Seurat` HVG selection. We found that many high-expression genes did not get selected by this method, including known marker genes like *Snap25* (Supplementary Figure S8a). The t-SNE embedding based on the default `Scanpy` preprocessing was similar to ours, but arguably showed less local structure (Supplementary Figure S8b). In the absence of ground truth cell labels, it is impossible to assess the representation quality objectively; however, based on the variance of known marker genes, we argue that compound Pearson residuals provide a more meaningful representation of the data.

As noted above, computing compound Pearson residuals was fast. For this dataset with ca. 23 000 cells and 38 000 genes it took ∼15 s on a single CPU. The resulting dense matrix of residuals used 3.4 Gb of RAM instead of 1.6 Gb for the sparse matrix of read counts. Census and qUMI had slower runtimes (3 h and 3 min respectively).

### 2.5 Compound Pearson residuals recover ground truth

To confirm that compound Pearson residuals are indeed able to recover true marker genes and true cell classes, we simulated read count data with known ground truth based on the Tasic et al. (2018) dataset. In our simulations, sequencing depths *n*_*s*_, gene fractions *p*_*g*_, and class identities were taken from the real data, and we used a compound model with NB(*θ* = 100) as UMI distribution and a geometric distribution as amplification distribution to simulate counts within each class. The true *α*_*Z*_ in this case is equal to 199 (see Methods).

In *Simulation I*, we allowed only a small set of known marker genes to vary between classes. Compound Pearson residuals showed high residual variance only for those ground truth marker genes (even when *α*_*Z*_ was misspecified, Supplementary Figure S9a–b). In contrast, UMI Pearson residuals (*α*_*Z*_ = 1) showed high residual variance for many non-variable genes (Supplementary Figure S9c).

In *Simulation II*, we mirrored the full cluster structure of the data by using cluster-specific *p*_*g*_ values for all genes. Using compound Pearson residuals, we obtained reasonable residual variances and embeddings recovering ground truth clusters (regardless of the exact value of *α* ≫ 1, Supplementary Figure S9d–e, g–h). At the same time, UMI Pearson residuals failed to stabilize the variance and incorrectly merged many clusters (Supplementary Figure S9f,i). In summary, both simulation experiments confirmed that compound Pearson residuals can recover true marker genes and cell types.

### 2.6 The broken zeta distribution as amplification model

So far, we did not specify the amplification distribution Z. Instead, we only characterized its mean and variance through the *α*_*Z*_ parameter. While this was sufficient to compute compound Pearson residuals, an explicit amplification distribution is needed for the complete specification of the compound model, enabling likelihood calculation or using it as a generative model. To find an appropriate statistical model, we obtained empirical amplification distributions from experimental data generated with several UMI-based protocols: CEL-seq2, Drop-seq, MARS-seq, and SCRB-seq (Ziegenhain et al., 2017), as well as Smart-Seq3 (Hagemann-Jensen et al., 2020) and Smart-Seq3xpress (Hagemann-Jensen et al., 2022). Together, we analyzed 106.3 million UMIs from 856 cells.

For each UMI barcode, we computed the number of times it occurred in the sequenced reads (its copy number). The normalized histogram of these copy numbers provided an empirical characterization of the amplification distribution Z (Figure 4). Across all sequencing protocols, the copy number histograms in log-log coordinates showed a characteristic elbow shape: Higher copy numbers were less frequent, and the distribution followed two separate decreasing trends in two ranges of copy numbers. This shape can be described by a broken power law, i.e. two separate power laws for low and high copy numbers. The exact shape of the distribution differed between sequencing protocols, leading to different values of mean and variance (Table 1). These values are influenced by how deeply a sample is sequenced and how many cycles of amplification are employed: e.g. the Smart-seq3 Xpress dataset was sequenced shallower than the other two Smart-seq3 datasets (ca. 94 000 vs ca. 572 000–806 000 reads per cell) and with fewer PCR cycles, leading to substantially lower 𝔼 [*Z*] values.

**Table 1:**
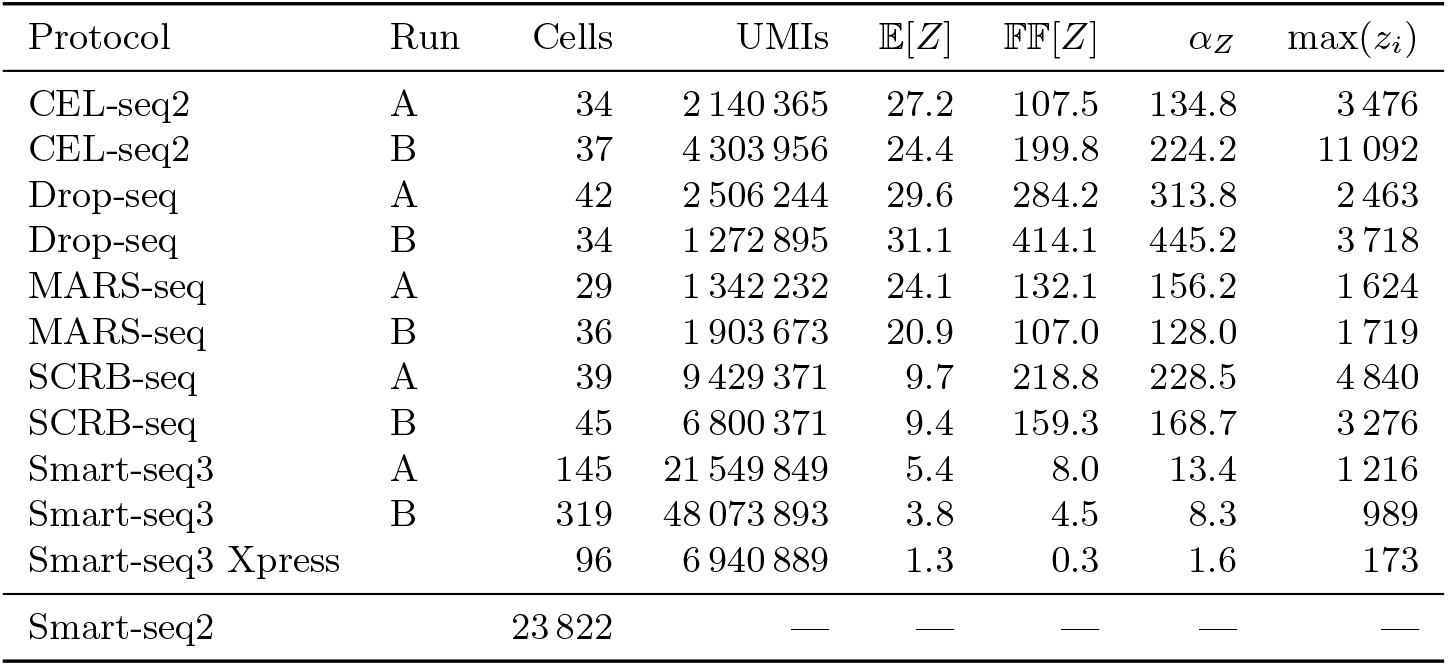
Key statistics of the observed amplification distribution across protocols. Top part: Each row in the table corresponds to one of the datasets presented in Figure 4. The number of UMIs shows how many observed copy numbers *z*_*i*_ were used to compute the statistics for that dataset. Bottom row: The Smart-seq2 dataset analyzed in Figure 3, where UMIs were not observed.

**Figure 4:**
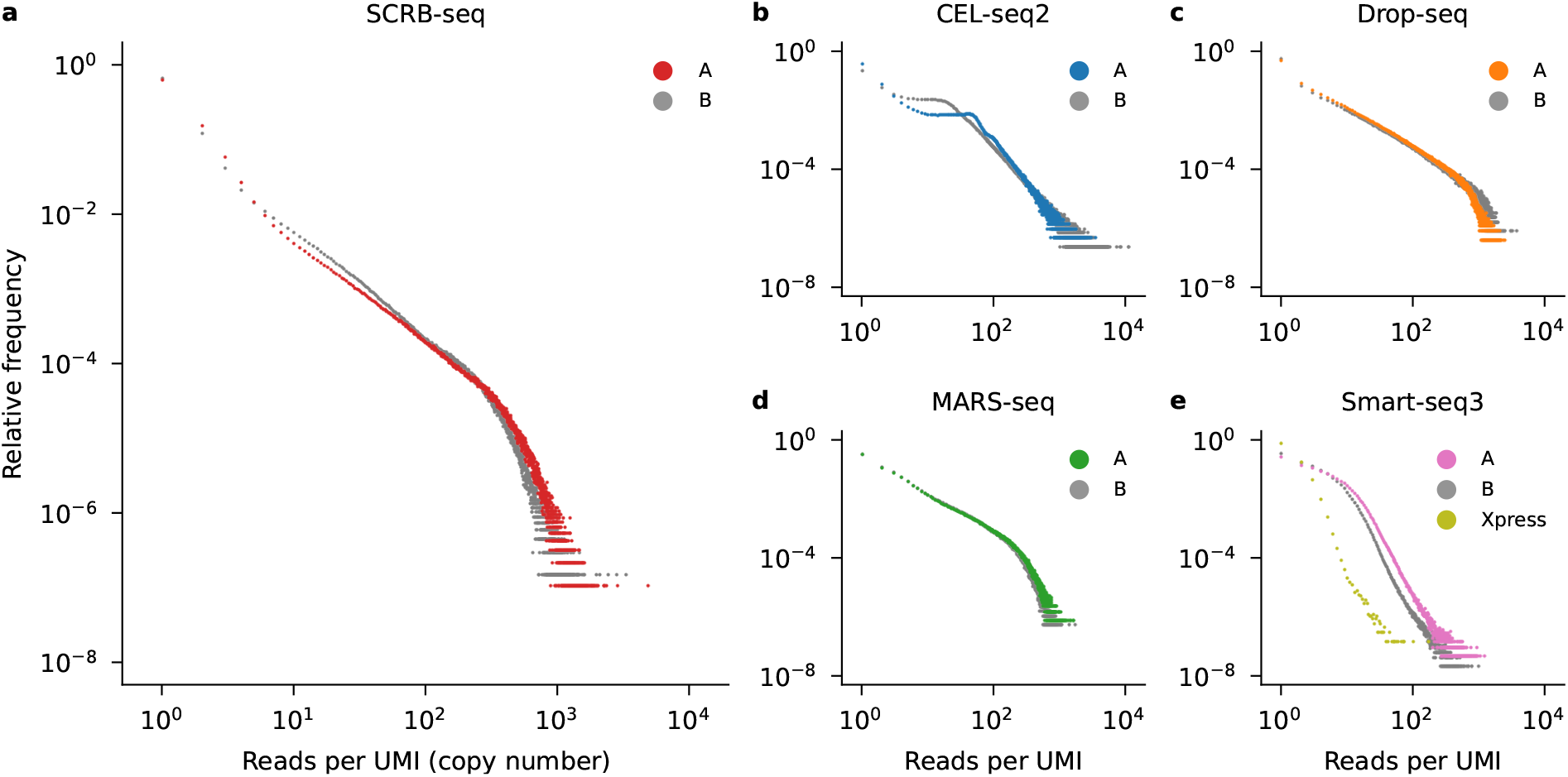
Observed amplification distributions follow a similar shape across protocols. Each panel shows a distribution of UMI copy numbers for a given UMI protocol. **a–d:** SCRB-seq, CEL-seq2, Drop-seq, and MARS-seq data from Ziegenhain et al. (2017). For each protocol two identical runs were performed (A and B). **e:** Smart-seq3 protocols. Data from a single-end experiment (A), a paired-end experiment (B) (Hagemann-Jensen et al., 2020), and a Smart-seq3 Xpress experiment (Hagemann-Jensen et al., 2022).

To study how stable the amplification distribution was across cells in the same sample, we computed per-cell estimates of 𝔼 [*Z*] and *α*_*Z*_ = 𝔼 [*Z*] + 𝔽 𝔽 [*Z*]. The estimates showed some variability across cells (Supplementary Figure S10), but it was small enough that our assumption of the shared amplification parameters seems justified in practice. Moreover, the per-cell estimates of *α*_*Z*_ were correlated with the total number of read counts per cell (Supplementary Figure S11), and this between-cell variability is accounted for in our model in any case.

Note that for all protocols, the empirical distribution of copy numbers was monotonically decreasing, meaning that *z* = 1 was the most likely copy number, despite many cycles of amplification. This may seem counter-intuitive, but previous work (Best et al., 2015) showed that a mechanistic model of the amplification process followed by Poisson sampling at the sequencing stage can give rise to similar copy number histograms with power law behaviour.

As copy numbers are positive integers, we modeled the distribution of copy numbers *z* with a discrete broken power law. The discrete probability distribution with the mass function following a power law is called zeta distribution. For the broken power law, we adopted the term *broken zeta distribution*, which we define as having the following probability mass function (PMF):

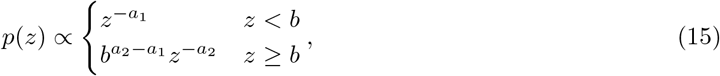

where *a*_1_ *>* 0 and *a*_2_ *>* 0 are negative slopes of the PMF in log-log coordinates, and *b* ∈ ℕ is the breakpoint between the two slopes. (Figure 5a, inset). We could choose the values for these three parameters such that the broken zeta distribution approximately matched the observed copy number histograms. For example, we obtained a good match for the Drop-seq protocol using *a*_1_ = 1.4, *a*_2_ = 4.5, and *b* = 500 (Figure 5a).

**Figure 5:**
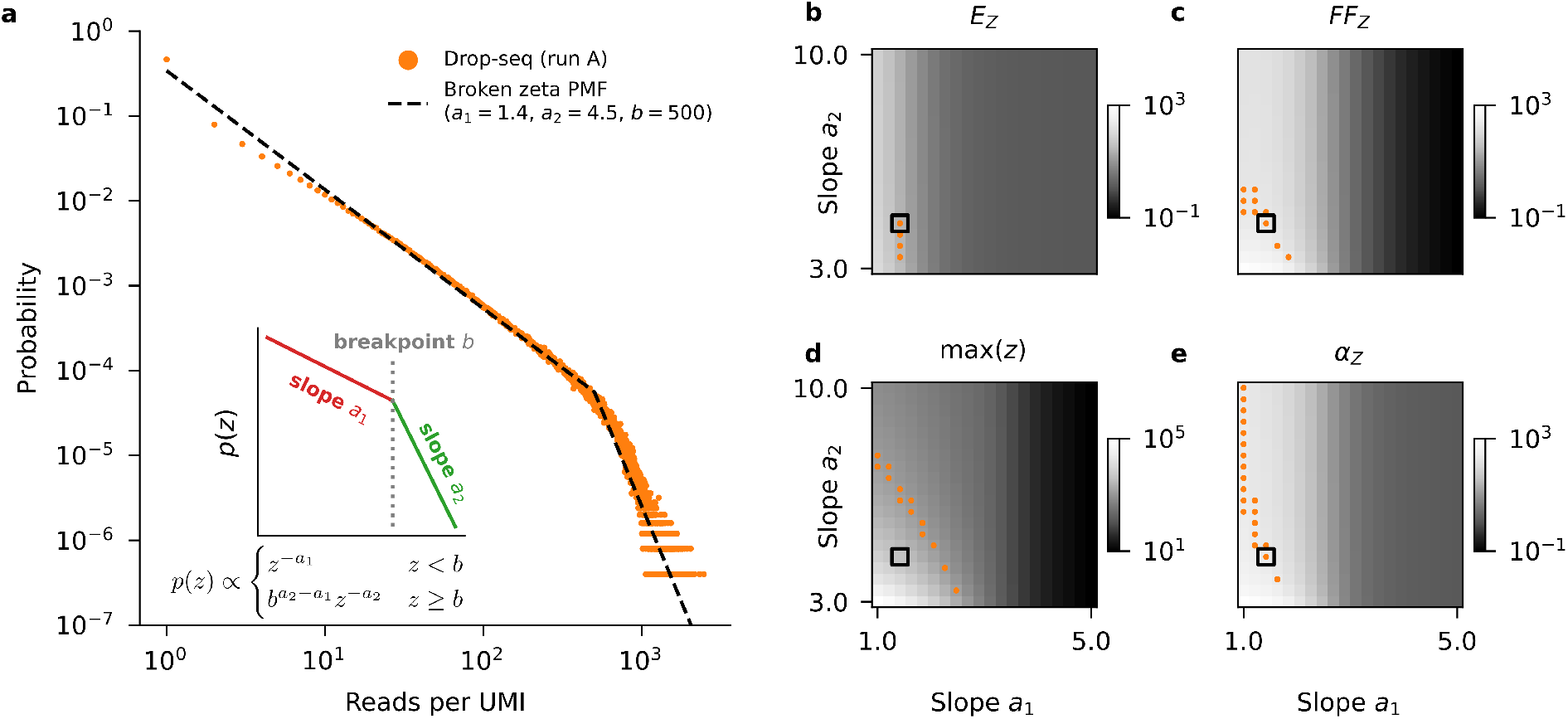
Broken zeta model can fit observed amplification distribution. **a:** Observed amplification distribution for Drop-seq (same as in Figure 4c) (orange dots) and the PMF of a broken zeta model (black line). The inset illustrates the parameters of the broken zeta model. **b:** The heatmap shows how the two slope parameters affect the mean of the broken zeta distribution. For each combination of *a*_1_ and *a*_2_, we sampled from a broken zeta model with these slope parameters and fixed *b* = 500. We used the same sample size as in the observed data in (a). The orange dots highlight the simulations yielding the sample mean close the observed mean (*±*10%). The black square shows the simulation corresponding to the fit shown in (a). **c–e:** Same as (b), but showing the sample Fano factor, sample maximum, and sample *α*_*Z*_.

The fitted model could reproduce several key statistics of the experimental data, such as the mean, the variance, and the Fano factor (Figure 5b–e). However, the broken zeta distribution produced sample maxima that were larger than empirically observed maxima (given the same sample size) (Figure 5d). This is a limitation of the broken zeta model as it tends to allocate non-zero probability mass to very high copy numbers that are not observed in practice. A more flexible model that limits the probability of very large copy numbers could potentially fit the data even better, but we considered the broken zeta distribution sufficient for our purposes.

### 2.7 Compound NB model with broken zeta amplification captures trends in read count data

The compound NB model (Equations 5–8) together with the broken zeta amplification distribution (Equation 15) provides a generative probabilistic model of the read counts in a biologically homogeneous population. To confirm that the model gives rise to realistic data, we used it to sample read counts and compared them to observed read count histograms in a biologically homogeneous dataset (Figure 6). We used the same dataset as above in Figure 2. For the amplification distribution, we used broken zeta parameter settings (*a*_1_ = 0.36, *a*_2_ = 5.1, *b* = 56) that led to an amplification model with *α*_*Z*_ = 50 and 𝔼 [*Z*] = 30, as we showed earlier that these values fit the protein-coding genes in this dataset well (Figure 2).

**Figure 6:**
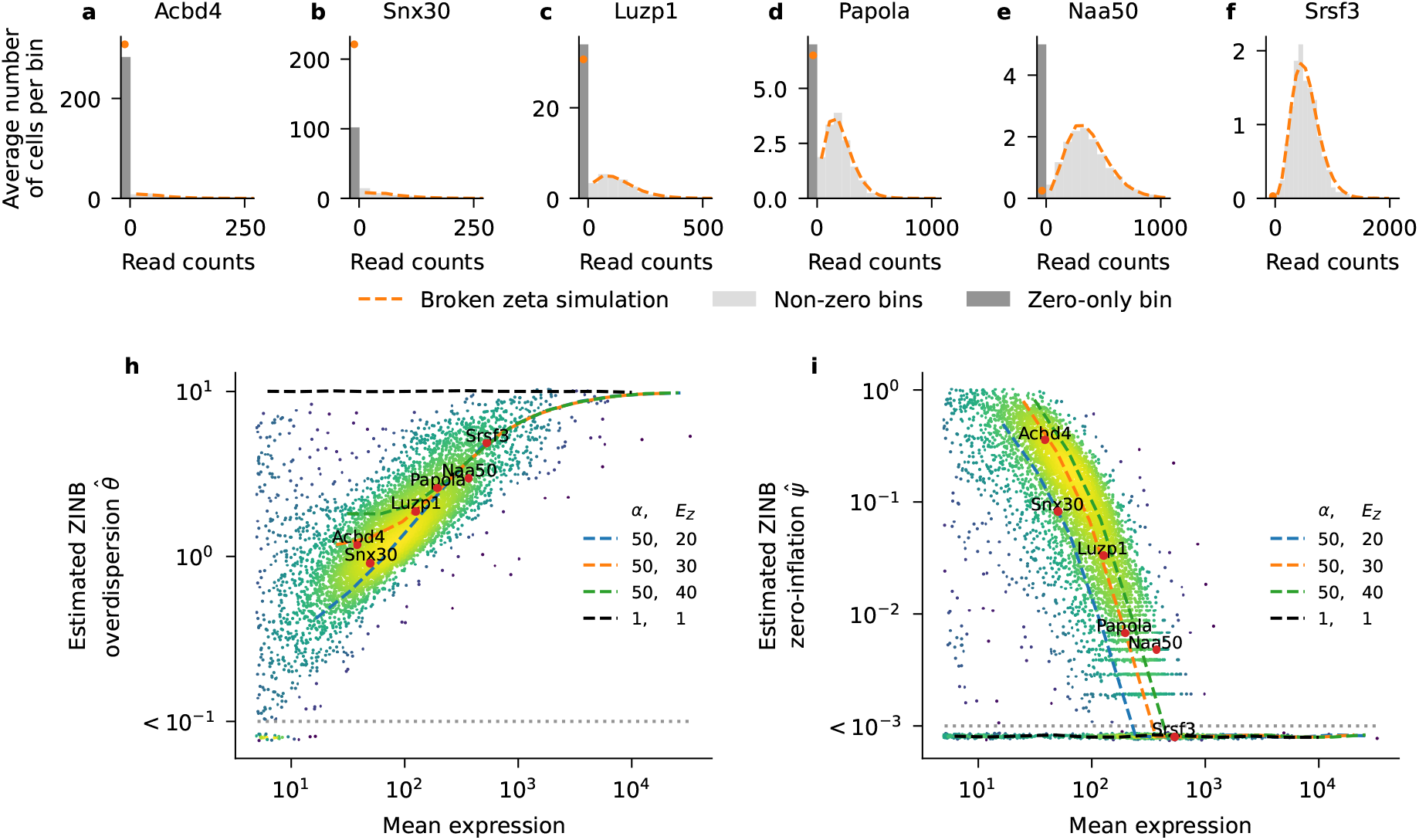
Broken zeta compound model simulations reproduce trends in read counts. Using a homogeneous subset (*n* = 1 049) of the mouse cortex data (from Figure 2), we fitted a zero-inflated negative binomial (ZINB) distribution to each gene individually. We also used the broken zeta model (*a*_1_ = 0.36, *a*_2_ = 5.1, *b* = 56 to sample read counts with a given mean expression. **a–f:** Each panel shows the observed read counts for a certain gene (gray histogram) and the histogram of counts sampled from the broken zeta model (orange line). Exact zeros are shown in a separate bin (orange dots, dark-gray bar). To show the zero-bin (width 1) and non-zero bins (width ≫ 1) on the same scale, the *y*-axis shows the average count per bin. Genes are ordered from left to right by mean expression. Note that with higher expression, fraction of zeros decreases, and the histogram shape changes from a geometric-looking distribution to a Poisson-looking distribution. **h:** Estimated ZINB overdispersion parameter 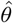 as a function of mean expression. Each dot is a gene, colored by local density of points. Values 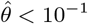 were clipped. Only genes with mean expression ≥ 5 are shown. Colored lines show 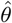 for samples from four different broken zeta models. For each model, we sampled counts over a range of expression fractions *p*_*g*_ for 10^5^ cells, each with fixed sequencing depth of *n*_*c*_ = 100 000. We used overdispersion *θ* = 10 for all genes. Broken zeta parameters: see Table 2. The black line corresponds to a negative binomial distribution (UMI model without amplification). Red dots highlight genes from panels (a)–(f). **i:** Estimated ZINB zero-inflation parameter 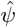 as a function of mean expression. Otherwise same as (h); values 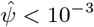 were clipped.

We found that the empirical count distributions of real genes (Figure 6a–f, grey) could be well matched by the compound NB model (Figure 6a–f, orange). Note that there is only one free parameter per gene in our simulation: *p*_*g*_, the fraction of RNA molecules taken up by this gene. The entire mass function is then determined by this single parameter. While the compound model did not fit every example gene perfectly (Figure 6b,e), it correctly captured the shapes of the distributions. In particular, for low-expression genes, the compound model predicted strong zero inflation and monotonically decreasing probability of non-zero counts (Figure 6a–b), while predicting a bell-shaped distribution without excess zeros for high expression genes (Figure 6f).

**Table 2:** Broken zeta parameters used for compound model read count simulations. Each row in the table corresponds to one of the four compound model simulations shown in Figure 6. Observed values refer to the obtained sample mean and sample variance (*n* ≈ 2 billion, see text for simulation details). The model with *α*_*Z*_ = 1 corresponds to the UMI case with constant copy number *z*_*i*_ = 1, and thus has no broken zeta parameters.

| $\mathbb{E}[Z]$ | | $\alpha_Z$ | | $a_1$ | $a_2$ | $b$ |
| --- | --- | --- | --- | --- | --- | --- |
| target | observed | target | observed |  |  |  |
| 1 | 1.0 | 1 | 1.0 | — | — | — |
| 20 | 20.1 | 50 | 50.4 | 0.96 | 15.1 | 91 |
| 30 | 30.2 | 50 | 51.4 | 0.36 | 5.1 | 56 |
| 40 | 37.9 | 50 | 50.5 | 0.01 | 17.6 | 71 |

In order to study these patterns more systematically, we fitted a zero-inflated negative binomial (ZINB) distribution to the count histograms of each gene separately. ZINB models have been used to model read counts before (Lopez et al., 2018), as read count data commonly exhibit zero-inflation compared to NB (Cao et al., 2021; Chen et al., 2018). A ZINB distribution has three parameters: the mean *µ*, the inverse overdispersion *θ*, and the zero-inflation parameter *ψ*. Its mass function is simply a negative binomial mass function with additional mass *ψ* on zero:

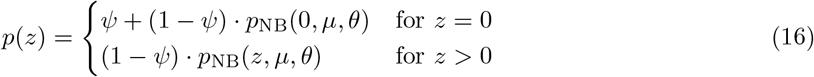

where *p*_NB_ is the NB probability mass function (see Methods). The ZINB distribution reduces to the NB distribution when *ψ* = 0. As in NB, the overdispersion parameter *θ* controls the shape of the distribution: *θ* = 1 corresponds to the geometric distribution with monotonically decreasing *p*(*x*), while higher values of *θ* result in more Poisson-like bell shapes (*θ* = ∞ corresponds to the Poisson case).

By fitting the ZINB model, we obtained independent estimates 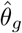 and 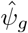 for each gene (Figure 6h–i). These estimates exhibited the same two patterns illustrated above for single genes. First, with increasing mean expression, genes tended to have a higher 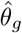, corresponding to smaller variance, and transitioned from a geometric-like to a Poisson-like shape. Second, with increasing mean expression, genes tended to have a lower 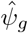, corresponding to less pronounced zero inflation.

The ZINB model cannot explain these trends, as all parameters *θ*_*g*_ and *ψ*_*g*_ can be chosen independently. In contrast, our compound model naturally gives rise to both effects. To demonstrate this, we repeated the ZINB fitting procedure on counts sampled from various compound NB models (see Methods for the broken zeta parameters), and reproduced both observations over a wide range of mean expressions (Figure 6h–i, colored lines). As expected, the model matching this dataset’s amplification parameters (*α*_*Z*_ = 50 and 𝔼 [*Z*] = 30, orange line, cf. Figure 2) provided the best match to the bulk of the distribution.

As a sanity check, sampling read counts from a NB model without amplification (*α*_*Z*_ = 1) and fitting ZINB distribution to the resulting samples recovered the original parameters (Figure 6h–i, black lines): constant overdispersion *θ* = 10 and absent zero inflation *ψ* = 0. This again shows that a NB model without amplification cannot describe the properties of the read count data. However, our results suggest that it is not necessary to include explicit zero-inflation like in a ZINB model, as it is naturally arising through the compound model.

## 3 Discussion

In this paper, we derived a parsimonious and theoretically grounded statistical model describing scRNA-seq read count data without UMIs. Furthermore, we showed that our compound model leads to analytic compound Pearson residuals, a fast, simple, and effective normalization approach for non-UMI data.

Despite the popularity of UMI protocols (Svensson et al., 2020), full-length non-UMI protocols such as Smart-seq2 (Picelli et al., 2013) remain relevant as they have higher sensitivity (Ziegenhain et al., 2017; Ding et al., 2020) and allow quantification of reads over full transcripts. This makes read count data indispensable for detection of splicing variants (Feng et al., 2021) or profiling of complex tissues with rare cell types (Tasic et al., 2018; Yao et al., 2021). The recently developed Smart-seq3/Smart-seq3xpress protocols (Hagemann-Jensen et al., 2020, 2022) contain UMIs on the 5’-end reads but do not have UMIs on internal reads, so our treatment remains relevant for Smart-seq3/Smart-seq3xpress as well.

While UMI counts can be modeled by a Poisson or a negative binomial (NB) distribution (Grün et al., 2014; Chen et al., 2018; Hafemeister and Satija, 2019; Townes et al., 2019; Svensson, 2020; Grün, 2020; Sarkar and Stephens, 2021; Rosales-Alvarez et al., 2023; Neufeld et al., 2023), read counts can not (Chen et al., 2018; Cao et al., 2021). Instead, they are often modeled by a more flexible zero-inflated negative binomial distribution (ZINB) (Pierson and Yau, 2015; Zappia et al., 2017; Chen et al., 2018; Risso et al., 2018; Lopez et al., 2018). However, this leaves unexplained what causes zero inflation and why there are relationships between the gene-specific ZINB parameters, such as less zero inflation for higher mean expression (Figure 6i).

Our compound model answers these questions. We showed that read counts in biologically homogeneous data can be well described by a compound negative binomial distribution, arising from simple statistical assumptions about the amplification and sequencing processes. Furthermore, we showed empirically that the distribution of copy numbers approximately follows a broken zeta distribution. Together, our compound NB model with amplification modeled by broken zeta yields a generative model reproducing zero-inflation and overdispersion patterns similar to what is observed in read count data. Compared to the ZINB model with three per-gene parameters (Equation 16), our model contains only one free per-gene parameter (Equations 5–8), and the varying zero-inflation and overdispersion naturally emerge as a function of a gene’s mean expression.

We observed that the distribution of copy numbers in UMI-containing data followed a similar shape across various protocols (Figure 4), implying that this is a general property of scRNA-seq data. We argued that this shape could be described by a broken power law, and hence we modelled it with a broken zeta distribution. This model is phenomenological, but previous work on mechanistic modelling of PCR amplification followed by Poisson sampling showed that these processes can give rise to similar copy number distributions (Best et al., 2015). We note that fitting parameters for power-law-like data is intrinsically difficult: common approaches such as least squares often return unstable estimates due to low-probability events (Clauset et al., 2009), which is why we avoided automatic parameter fitting. Instead, we qualitatively showed that the broken zeta model can give rise to realistic read count distributions.

Our compound NB model did not describe all genes perfectly: we found that a subset of genes, mostly pseudogenes, did not follow the compound model, but rather behaved as if they were not amplified (Figures 2 and 3). Similar bimodal patterns in gene variance have been be observed in previous works (e.g. Brennecke et al., 2013; Ziegenhain et al., 2017). Pseudogenes are copies of functional genes that contain a mutation making the copy dysfunctional. We can only speculate about the reason causing pseudogene read counts to have less variance: they may behave differently during amplification or sequencing, or perhaps their counts are an artifact of the alignment algorithm (all datasets we analyzed used STAR (Dobin et al., 2013)). In practice, such pseudogenes have less variance than expected under the compound NB model, so will be filtered out by the gene selection step in our suggested workflow (Figure 3a).

On the practical side, we used the compound NB model to derive a fast and theory-based normalization procedure for read counts: compound Pearson residuals. They constitute an extension of the UMI Pearson residuals normalization, that has proven to be effective for gene selection and normalization of UMI data (Hafemeister and Satija, 2019; Lause et al., 2021). We showed that compound Pearson residuals work well for processing complex read count datasets, leading to a biologically meaningful gene selection and embeddings. Importantly, we also showed that normalization and gene selection using the non-compound UMI Pearson residuals leads to suboptimal results on read count data, underscoring the importance of an adequate statistical model.

Compound Pearson residuals only require to set the *α*_*Z*_ parameter of the amplification distribution. Whereas *α*_*Z*_ can be observed directly from the copy number distribution for UMI-containing data (i.e. reads per UMI, Table 1), it is unknown in a Smart-seq2 experiment. Reassuringly, we found that the results of compound Pearson residuals do not strongly depend on the exact value of *α*_*Z*_ — as long as it is set within a reasonable range ≫ 1, such as *α*_*Z*_ ∈ [10, 1000]. When working with Smart-seq2 data, we recommend using *α*_*Z*_ = 50 by default. Furthermore, it is possible to empirically adjust *α*_*Z*_ to a given dataset from any sequencing protocol. Indeed, under the common assumption that most genes are not differentially expressed, the majority of genes should have residual variance close to one. Thus, adjusting *α*_*Z*_ until this condition is fulfilled will typically lead to a reasonable setting (Supplementary Figure S6).

Typical approaches to read count data normalization consist of scaling read counts by a size factor to account for sequencing depth (CPM: counts per million) and sometimes gene length (TPM: transcripts per million (Li and Dewey, 2011), or RPKM: reads per kilobase per million (Mortazavi et al., 2008)), followed by a log-transform (Luecken and Theis, 2019; Andrews et al., 2021; Slovin et al., 2021). Various methods have been suggested to estimate the required size factors, going beyond CPM/TPM/RPKM (Vallejos et al., 2017): via spike-ins (Brennecke et al., 2013; Lun et al., 2017), cell pooling (Lun et al., 2016), housekeeping genes (Andrews et al., 2021), separate scaling for groups of genes (Bacher et al., 2017), or a Bayesian approach (Tang et al., 2020). However, all of these methods depend on the log-transform for variance stabilization, which is inherently limited (Lun, 2018) and fails to fully stabilize the variance (Ahlmann-Eltze and Huber, 2023). In contrast, our compound Pearson residuals use the mean-variance relationship that follows from simple statistical assumptions, and the resulting residuals are variance-stabilized by design and do not require any explicit normalization by the gene length.

Two existing methods aim to transform read counts so that their distribution matches the distribution of UMI counts. Census (Qiu et al., 2017) linearly scales the read counts within each cell to set the mode of the count distribution to 1, while qUMI (Townes and Irizarry, 2020) performs quantile normalization within each cell to transform the entire distribution to the typical shape of within-cell UMI counts. In both cases, the transformations are heuristics not based on any generative statistical model, and the transformed data still require UMI-specific normalization. In contrast, our compound Pearson residuals perform necessary normalization directly on the read counts. In practice, we observed that qUMI and Census lead to comparable normalization results as our compound Pearson residuals, but our method follows from an explicit statistical model that offers theoretical insights into the data generation process underlying read counts. For example, as described above, our model captures previously unexplained patterns in the zero-inflation and overdispersion of read count data.

One limitation of our model is that it assumes that the amplification distribution is the same for all genes and cells and uses the single amplification parameter *α*_*Z*_ shared by all cells. This is not the case in Census and qUMI, which both use cell-specific adjustments. Reassuringly, we did not observe strong cell-to-cell variability in *α*_*Z*_ estimates (Supplementary Figure S10), and furthermore found that *α*_*Z*_ correlated with total counts per cell (Supplementary Figure S11) — a factor which our model explicitly accounts for.

In summary, we show that the compound NB distribution is the appropriate statistical model for read count data, naturally giving rise to compound Pearson residuals as an effective, convenient and theoretically motivated way of data pre-processing.

## 4 Methods

### 4.1 Datasets and preprocessing

Our example read count dataset throughout this paper is the mouse brain dataset from Tasic et al. (2018), GEO accession GSE115746. It contains cells from the primary visual cortex (VISp) and the anterior lateral motor area (ALM), and was sequenced with Smart-seq2. We downloaded the data for both areas from https://portal.brain-map.org/atlases-and-data/rnaseq/mouse-v1-and-alm-smart-seq and used only the exonic counts and applied the same cell filtering as Tasic et al. (2018), leading to 23 822 cells and 42 776 genes with at least one count. We used the Python package mygene to query the `mygene.info` database (Wu et al., 2013) for gene type annotations (`type of gene field`) with the Entrez gene identifiers. We queried `BioMart` to obtain transcript lengths for all Ensemble mouse genes (database *GRCm39*, column ‘Transcript length (including UTRs and CDS)’, Cunningham et al. (2022)).

From these data, we assembled a biologically homogeneous subset by selecting only cells from one of the largest neuronal clusters (named *VISp Penk Col27a1* by Tasic et al. (2018), 1 049 cells, 33 914 detected genes). These data were used without further filtering in Figures 2 and 6 and Supplementary Figures S1 and S2.

To assemble a heterogeneous dataset, we took the full dataset but filtered out genes that were detected in less than 5 cells as in previous work on UMI Pearson residuals (Hafemeister and Satija, 2019; Lause et al., 2021), leading to 23 822 cells and 38 510 genes.

To study copy number distributions across protocols, we used the following UMI datasets:

- Mouse embryonic stem cells profiled by CEL-seq2, Drop-seq, MARS-seq, and SCRB-seq (Ziegenhain et al., 2017); GEO accession GSE75790;
- Mouse fibroblasts profiled with Smart-seq3 paired-end (Johnsson et al., 2022); accession E-MTAB10148, sample `plate2`;
- Mouse fibroblasts profiled with Smart-seq3 single-end (Hagemann-Jensen et al., 2020); accession E-MTAB-8735, sample `Smartseq3.Fibroblasts.smFISH`;
- HEK293 cells profiled with Smart-seq3Xpress (Hagemann-Jensen et al., 2022); accession E-MTAB11467.

For UMI deduplication, we used Hamming distance correction with a threshold of 1. See Table 1 for numbers of cells and UMIs per dataset. The reads-per-UMI tables are available at https://zenodo.org/record/8172702.

We used scanpy 1.9.0 (Wolf et al., 2018) and `anndata` 0.8.0 (Virshup et al., 2021) for all scRNA data handling in `Python` 3.8.10, along with `sklearn` 1.0.2 (Pedregosa et al., 2011), `numpy` 1.21.5 (Harris et al., 2020), and matplotlib 3.5.1 (Hunter, 2007).

### 4.2 Simulation study

To generate a realistic validation dataset with ground truth marker genes *(Simulation I)*, we simulated read counts based on the Tasic et al. (2018) read counts *X*_*cg*_ as follows. For each gene *g*, we computed the average expression fractions *p*_*g*_ = Σ_*c*_ *X*_*cg*_*/ Σ*_*cg*_ *X*_*cg*_. For a set *G* of 14 well-known brain cell type marker genes (Tasic et al., 2016) and each of the 133 clusters in the Tasic et al. (2018) data, we computed the within-cluster fraction *p*_*ig*_, where *i* is the cluster index. For each cell *c*, we computed its total read counts *n*_*c*_ = Σ_*g*_ *X*_*cg*_ and assumed total UMI counts per cell 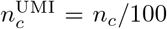. We then generated UMI counts as 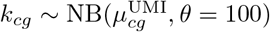, where

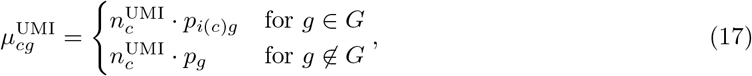

where *i*(*c*) denotes cluster assignments of cell *c*. In words, only the marker genes from *G* were allowed to differ between clusters. We then simulated the amplification of each UMI by drawing copy numbers from the shifted geometric distribution *z*_*i*_ ∼ Z = Geom_+_(*µ* = 100), which corresponds to amplification with 𝔼 [*Z*] = 100 and *α*_*Z*_ = 199 (see below). We finally summed the copy numbers for each gene and cell to obtain read counts 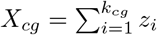 (Equation 5). After filtering out genes with less than 5 cells as above, *Simulation I* yielded 23 822 cells and 30 652 genes.

To obtain a second validation dataset with a richer cluster structure and ground truth cell types *(Simulation II)*, we used the same simulation setup as above, but allowed all genes to have cluster-specific fractions, i.e.

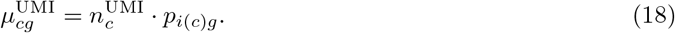

After filtering as above, *Simulation II* yielded 23 822 cells and 30 576 genes.

Both simulations generated copy numbers *z*_*i*_ from the shifted geometric distribution *Z* = Geom_+_(*µ* = 100), which is equivalent to *Z* = NB(*µ* = 99, *θ* = 1)+1, with *z*_*i*_ ∈ ℕ_+_ being positive integers. The variance of the negative binomial is equal to 99 + 99^2^*/*1 = 9900 and 𝔼 [*Z*] = 100, so the Fano factor is 99, leading to *α*_*Z*_ = 𝔼 [*Z*] + 𝔽 𝔽 [*Z*] = 199.

### 4.3 Mathematical details of the compound negative binomial model

We use the term *compound Poisson/NB distribution* to describe a discrete random variable that is constructed as a sum over a random number of i.i.d. terms. A compound model has an ‘inner’ and an ‘outer’ distribution: The inner distribution generates the i.i.d. summation terms (Equation 6), while the outer distribution governs the number of terms to be summed (Equation 7). This setup is known under various names: Johnson et al. (2005) uses the term *stopped-sum* distribution. When the outer distribution is the Poisson distribution, the compound model is known as compound Poisson (Adelson, 1966), stuttering Poisson (Kemp, 1967; Moothathu and Kumar, 1995), or generalized Poisson (Feller, 1943) distribution.

Note that the term *compound distribution* can also have a different meaning: for example, in their work on qUMI normalization, Townes and Irizarry (2020) used the term ‘compound Poisson model’ to describe a Poisson model with rate parameter *λ* governed by another distribution.

The expectation of a compound random variable 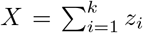 with inner distribution *z*_*i*_ ∼ *Z* and outer distribution *k* ∼ *K* can be obtained as follows:

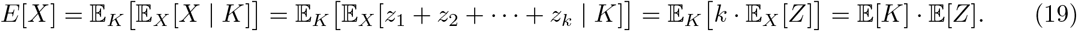

The variance can be computed similarly:

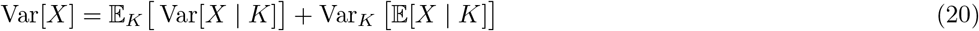

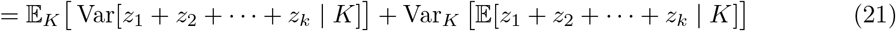

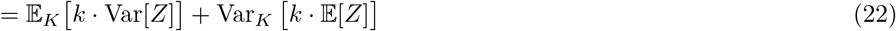

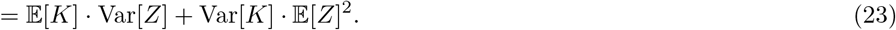

Together, this leads to the following mean-variance relationship:

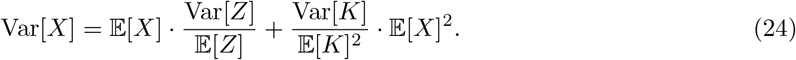

We use the negative binomial (NB) distribution as outer distribution in our compound model. The probability mass function for the NB distribution can be parametrized in several different ways. We use

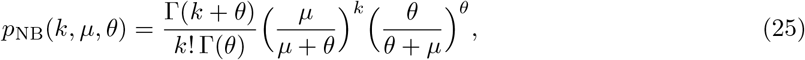

where *µ* is the mean and *θ* is the overdispersion parameter. The variance is then given by Var[*K*] = 𝔼 [*K*] + 𝔼 [*K*]^2^*/θ* = *µ* + *µ*^2^*/θ*.

Plugging the mean-variance relationship of *K* into the mean-variance relationship of *X*, we finally get

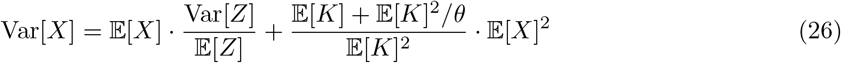

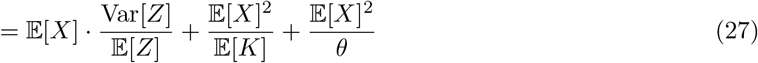

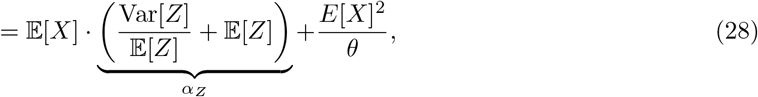

where we used the fact that 𝔼 [*X*] = 𝔼 𝔼*Z*] · E[*K*]. The relationship in Equation 28 yields the lines shown in Figure 2a.

From here we can obtain the relationship between the mean of *X* and the Fano factor of *X*:

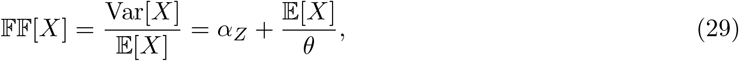

shown as lines in Figure 2b. For *E*[*X*] ≪ *θ*, this reduces to 𝔽 𝔽 [*X*] ≈ *α*_*Z*_.

To derive the relationship between the mean of *X* and the fraction of zero counts, we note that the inner distribution in our model is strictly positive (*z* ≥ 1). Any zero count *X* = 0 must thus originate from a *k* = 0 from the outer NB distribution. As a result, we can derive the fraction of zero counts from the NB probability mass function (Equation 25):

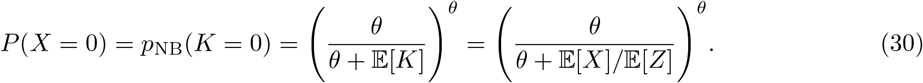

This relationship is shown as lines in Figure 2c. For *θ* → ∞, this converges to the Poisson case *P* (*X* = 0) = *e*^−𝔼 [*K*]^ = *e*^−𝔼 [*X*]*/* 𝔼 [*Z*]^.

### 4.4 Compound Pearson residuals

For gene selection with compound Pearson residuals, we computed 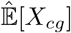 from the filtered r ead count matrix *X* (Equation 10) and then obtained residuals using Equation 13. We selected 3 000 highly variable genes (HVGs) with the highest residual variance. To normalize, we then subset the raw count data matrix to the HVGs, and computed compound Pearson residuals again on that subset, and used these re-computed residuals for further analysis (consistent with our previous work, Lause et al. (2021)). Using the residuals computed from the full data matrix and subsetting them to HVGs led to very similar results.

Unless otherwise stated, we used *α*_*Z*_ = 50 and *θ* = 100 for computing residuals, and clipped residuals to 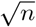 where *n* is the number of cells, following Hafemeister and Satija (2019) (see Lause et al. (2021) for a motivation for this heuristic).

### 4.5 Census counts and qUMIs

We obtained both Census counts and qUMIs via their official R implementations using `R 4.1.3`. To obtain Census counts, we used `bioconductor-monocle 2.22.0` (Huber et al., 2015; Qiu et al., 2017).

To obtain qUMIs, we used `quminorm 0.1.0` from http://github.com/willtownes/quminorm/ (Townes and Irizarry, 2020). As both methods expect TPMs as input, we subset the (Tasic et al., 2018) data to the 27 841 genes for which length annotations were available (see above), and computed TPM from read counts *X*_*cg*_ and gene lengths *l*_*g*_ (in kilobase) as

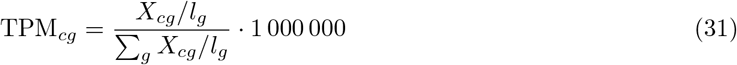

Running Census on the full matrix was very slow (*>*24 h), so we split the TPM matrix into batches of 1000 cells. This substantially sped up the computation. Filtering Census counts and qUMIs for genes with at least 5 cells yielded 23 822 cells and 25 248 genes.

### 4.6 t-SNE visualizations

As basis for all t-SNE embeddings, we computed the first 1 000 principal components (PCs) of the HVGs residuals with `sklearn 1.0.2` (Pedregosa et al., 2011). For all t-SNE embeddings, we used `openTSNE 0.6.0` (Poličar et al., 2019) with default settings unless otherwise stated. To ensure comparability between the t-SNE embeddings in Supplementary Figure S6, we used the first two PCs of the HVG residuals computed with *α*_*Z*_ = 10 (panel C) as shared initialization after scaling them with `openTSNE.initialization.rescale()`.

To visualize expression strength of a given gene across the t-SNE map(Figure 3c), we used squareroot-transformed depth-normalized counts

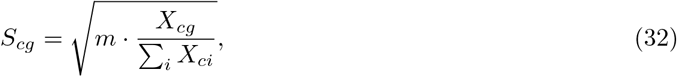

where *m* is the median row sum of *X*.

### 4.7 Fitting zero-inflated negative binomial (ZINB) models to single genes

To obtain per-gene estimates of overdispersion 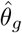 and zero inflation 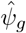 in the absence of biological variability, we fitted a ZINB model to the raw read counts of each gene in the *VISp Penk Col27a1* cluster (*n* = 1 049 cells). We used the `ZeroInflatedNegativeBinomialP.fit regularized()` function from `statsmodels 0.13.2` (Seabold and Perktold, 2010) with default parameters. Only the 11 549 genes with within-cluster mean expression *>*5 were included because low-expression genes suffered from unstable parameter estimates. All genes with fitting warnings (*n* = 2 064), fitting errors (*n* = 22) or invalid resulting estimates 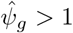 (*n* = 2 359) were excluded, such that 7 104 genes with valid converged estimates remained for further analysis and are shown in Figures 6h–i.

We applied the same fitting procedure to the simulated read counts shown as lines in Figure 6 (see below for simulation details). Here, 62 out of 100 simulated genes had a mean expression *>*5 and were used for fitting. 5 of them resulted in invalid 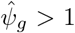 values and were excluded, such that 57 simulated genes remained for plotting and analysis.

### 4.8 Sampling copy numbers from the broken zeta model to simulate compound model read counts

The broken zeta model we describe in Equation 15 is the discrete version of the broken power law. A continuous probability distribution with probability density following a power law is called the Pareto distribution. Its discrete analogue is known under various names including Riemann zeta distribution (or simply zeta distribution), discrete Pareto distribution, and Zipf distribution(Johnson et al., 2005). We therefore refer to the discrete broken power law distribution as *broken zeta distribution*. While continuous broken power law distributions are commonly used in astrophysics (Jóhannesson et al., 2006), we are not aware of any prior use of discrete broken power law distributions for statistical modeling.

For the simulations in Figures 5 and 6, we sampled copy numbers from a given broken zeta distribution. For that, we computed the approximate PMF for a limited support *z* ∈ {1, 2, …, 10^5^} with Equation 15, and normalized the resulting probabilities to sum to 1. This way we did not need to compute the normalization constant in Equation 15.

For the simulations of the Drop-seq copy number distribution in Figure 5b–e we used *n* = 2 506 244 samples per parameter set, which is the number of UMIs in the Drop-seq A dataset. We extended the support until 10^6^ for this particular simulation, because we observed max(*z*_*i*_) ≈ 10^5^ for some of the more extreme parameter combinations.

In order to sample realistic read counts from four compound models with different combinations of mean copy number 𝔼 [*Z*] and *α*_*Z*_ (Figure 6), we used a grid search over the broken zeta parameters *a*_1_, *a*_2_, and *b* to find parameter combinations that best matched the required values. Table 2 lists the parameters and amplification statistics of the chosen models. Across all parameter sets shown in Figure 6, we observed max(*z*_*i*_) = 7 179 which was far below the end of the support.

We first sampled unique sequenced molecules *k*_*cg*_ from a negative binomial with *k*_*cg*_ ∼ NB(*p*_*g*_*n*_*c*_, *θ*) for 25 genes over a log-spaced range of expression fractions *p*_*g*_ ∈ [10^−8^, 10^−1^]. We did this for 10^5^ cells, each with fixed sequencing depth of *n*_*c*_ = 10^5^. We used overdispersion parameter *θ* = 10 for all genes. This led to a total of Σ*k*_*cg*_ = 2 047 196 087 simulated UMIs. Then, for each of them, we sampled a copy number *z*_*i*_ from the broken zeta model as described above, and summed over copy numbers for the same cell and gene to obtain a read count 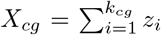. We used the same set of simulated UMIs for the four compound model simulations shown in Figure 6 and Table 2.

## Declarations

### Ethics approval and consent to participate

Not applicable.

### Consent for publication

Not applicable.

### Availability of data and materials

The datasets generated and/or analysed during the current study are publicly available as described in Methods (Section 4.1). All analysis code is available under the GNU General Public License v3.0 at https://github.com/berenslab/read-normalization, and is archived in the Zenodo repository https://zenodo.org/doi/10.5281/zenodo.12806891.

## Authors’ contributions

Conception of the method: JL, PB, DK. Data acquisition and preprocessing: CZ, LH. Data analysis and interpretation: JL, DK, CZ. Drafted the manuscript: JL. Revised the manuscript: All authors.

## Acknowledgements

We would like to thank Rickard Sandberg for discussions.

## Supplementary Figures

**Figure S1:**
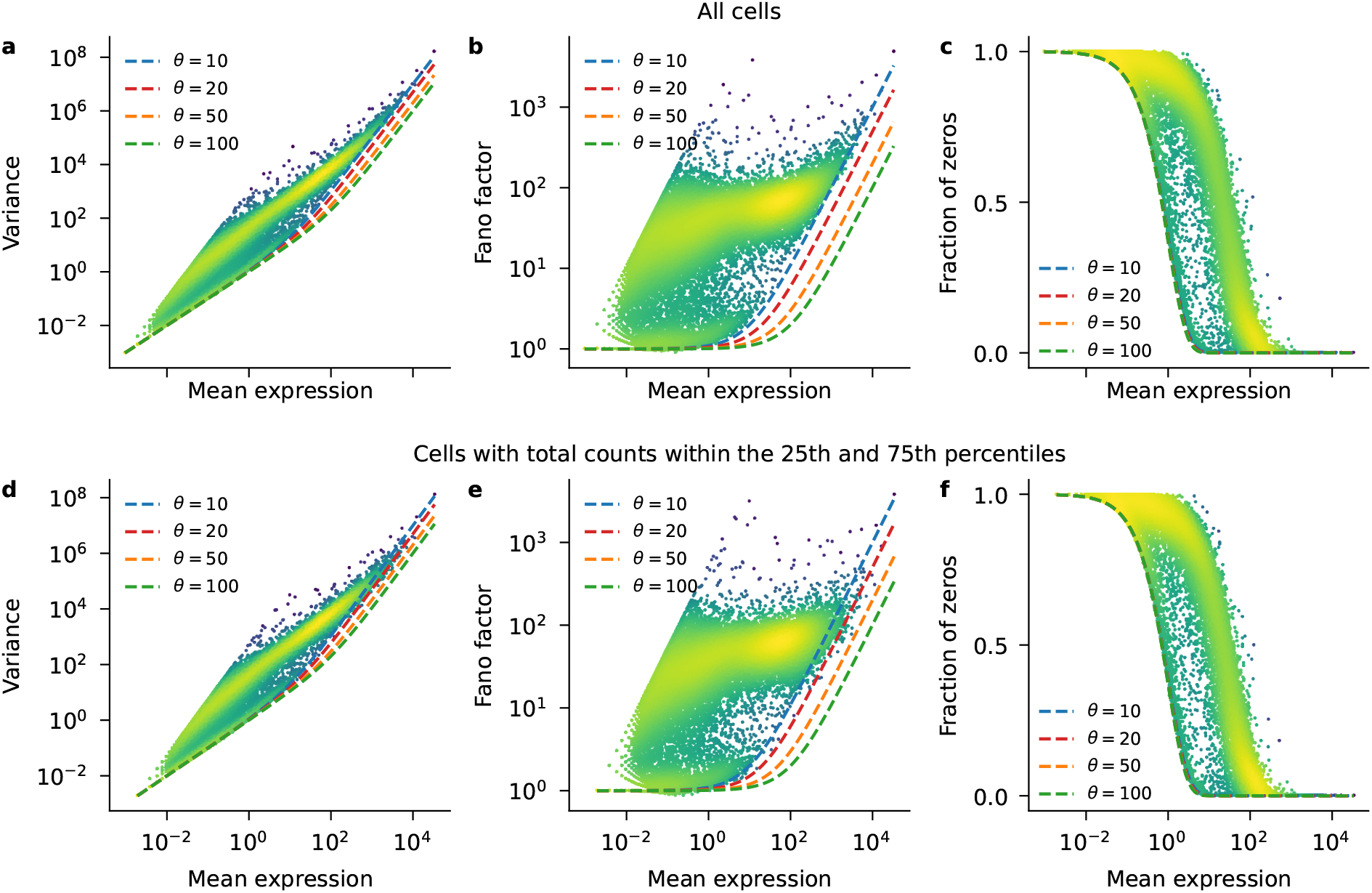
Sequencing depth variation increases the apparent overdispersion. Both rows are a reproduction of Figure 2, but showing pure NB models (*α*_*Z*_ = 1) that differ in their overdispersion parameter *θ*. **a–c:** Plots based on all 1 049 cells as shown in Figure 2 with total counts per cell ranging from ∼680 000 (1^st^ percentile) to ∼2.84 million (99^th^ percentile). Note that the NB model with *θ* = 10 fits the boundary of the data distributions. **d–f:** Plots based on a subset of the 523 cells with total counts within the 25th and 75th percentiles (from ∼1.54 million to ∼1.90 million reads). Now the NB model with *θ* = 20 fits the boundary of the data distributions better than with *θ* = 10. In other words, overdispersion is smaller when controlling for sequencing depth variation.

**Figure S2:**
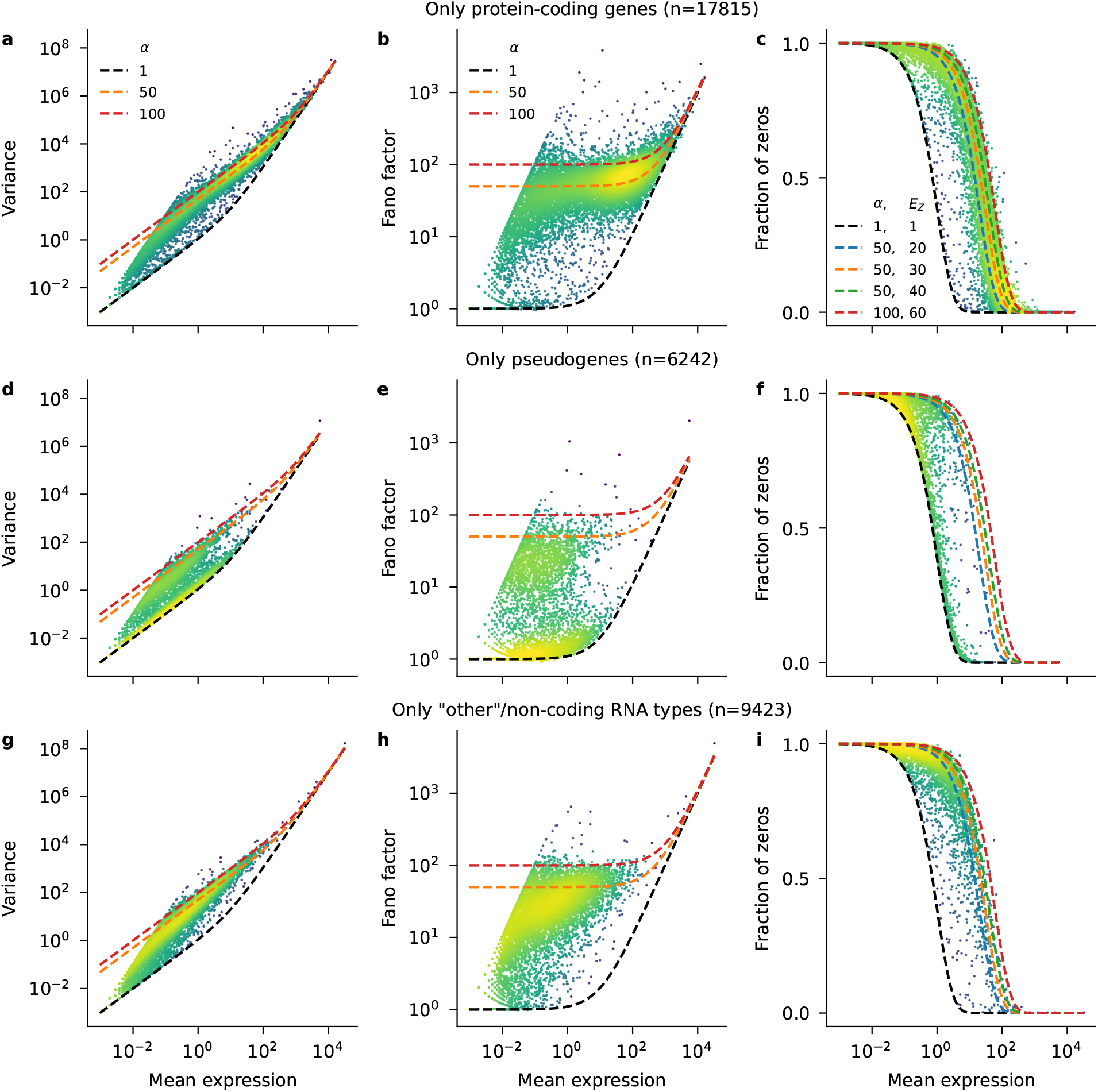
Non-amplified genes are mostly pseudogenes. Each row is a reproduction of Figure 2, but showing only genes from a certain category as obtained by the `mygene.info` annotation service. 434 genes without available annotation are not shown. **a–c:** Protein-coding genes. **d–f:** Pseudogenes. Many of them appear ‘non-amplified’ and do not follow the compound model, but rather the UMI model without amplification (black). **g–i:** Other, non-coding RNA species. Note that the bulk of these low-expression genes did not follow the compound model either.

**Figure S3:**
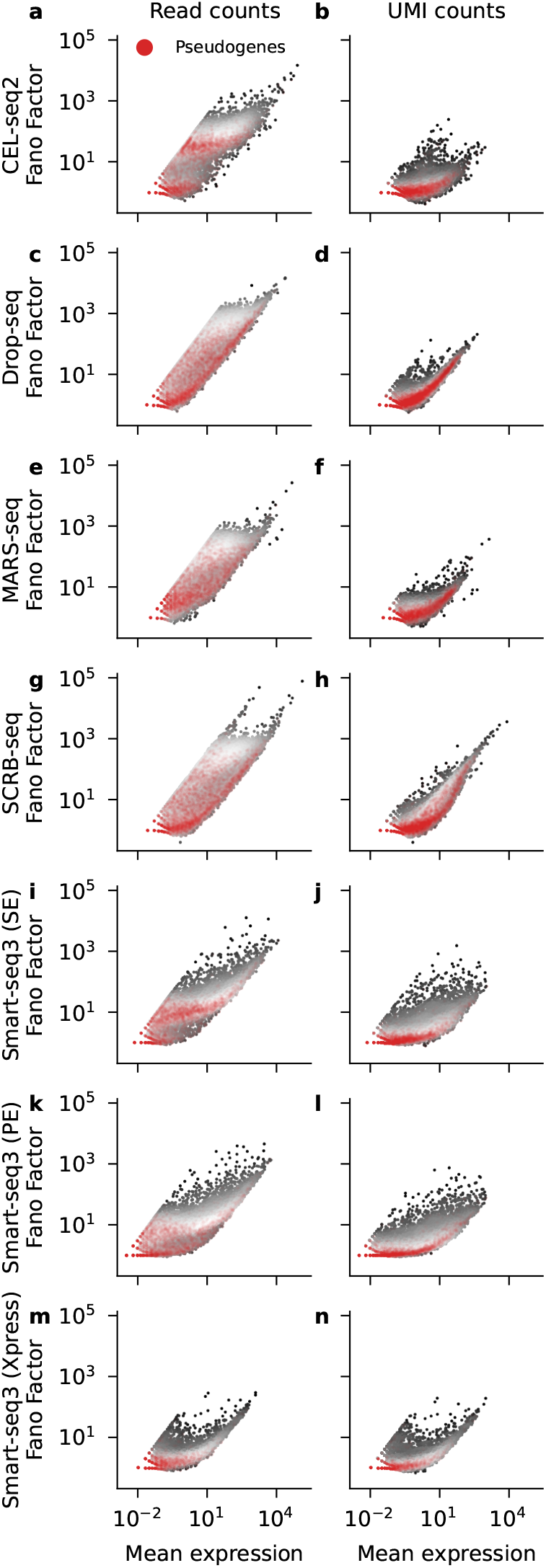
Pseudogenes have low Fano factor of read counts across protocols. Each panel shows the relationship between the mean and the Fano factor for all genes in each dataset. Higher density of dots is shown in brighter gray. Red dots with transparency show pseudogenes. Each row of panels shows a homogeneous dataset sequenced with a different protocol. All left-column panels are based on read counts, all right-column panels are based on UMI counts from the same dataset. Same data as in Figure 4. **a–h**: Mouse embryonic stem cells sequenced with various UMI protocols (Ziegenhain et al., 2017). For all protocols, only run A is shown. **i–l:** Smart-seq3 data from mouse fibroblasts (Hagemann-Jensen et al., 2020). SE: Single-end run. PE: Paired-end run. **m–n:** Smart-seq3 Xpress data from HEK293 cells (Hagemann-Jensen et al., 2022).

**Figure S4:**
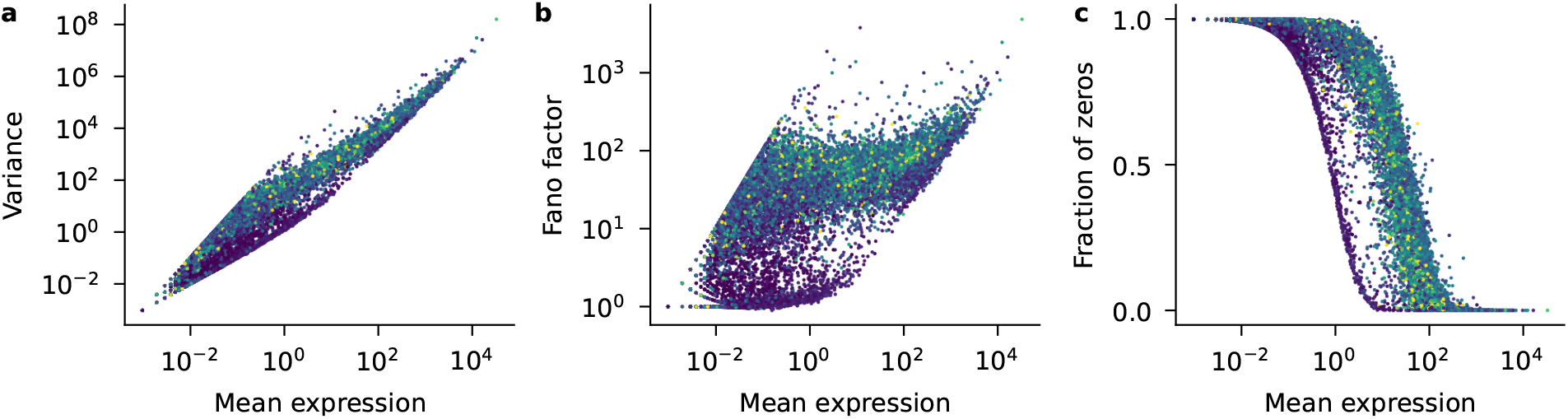
Pseudogenes have lower maximum transcript lengths. Reproduction of Figure 2, but showing each gene colored by the maximum length across all of its transcripts present in the Ensembl mouse gene database. 11 097 genes without transcript length annotation are not shown. Maximum lengths were clipped to the 98^th^ percentile (9 849 bp) before plotting.

**Figure S5:**
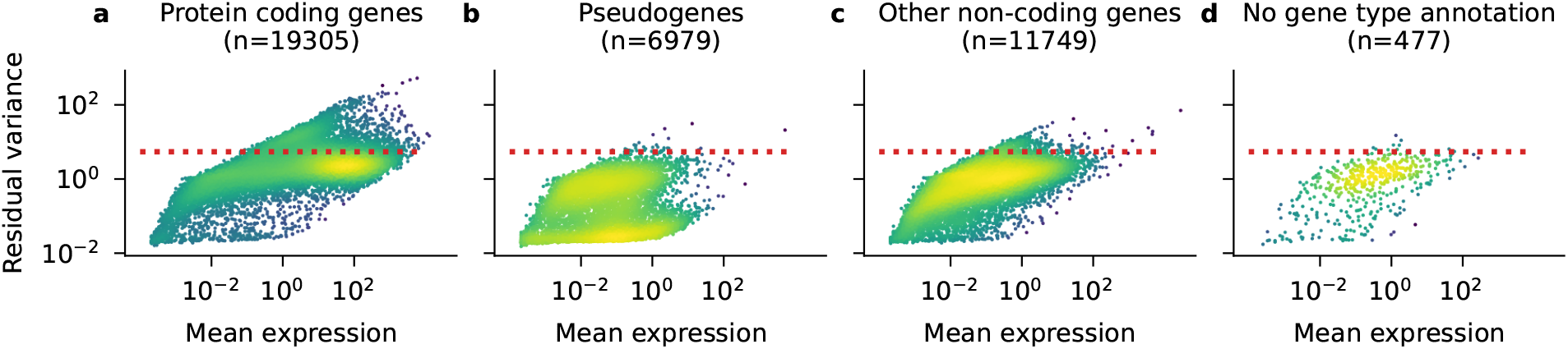
Genes with residual variance ≪ 1 are mostly pseudogenes. Each panel is a reproduction of Figure 3a, showing only a certain category of genes, as in Figure S2. Each dot represents a gene and shows its mean and residual variance in the full mouse visual cortex dataset (Tasic et al., 2018). Brighter color indicates higher density of points. Red line shows cutoff for selecting 3 000 HVGs among all genes. Gene type annotations taken from the `mygene.info` service. **a:** Protein-coding genes. **b:** Pseudogenes. **c:** Other, non-coding RNA species. **d:** Genes without available annotation.

**Figure S6:**
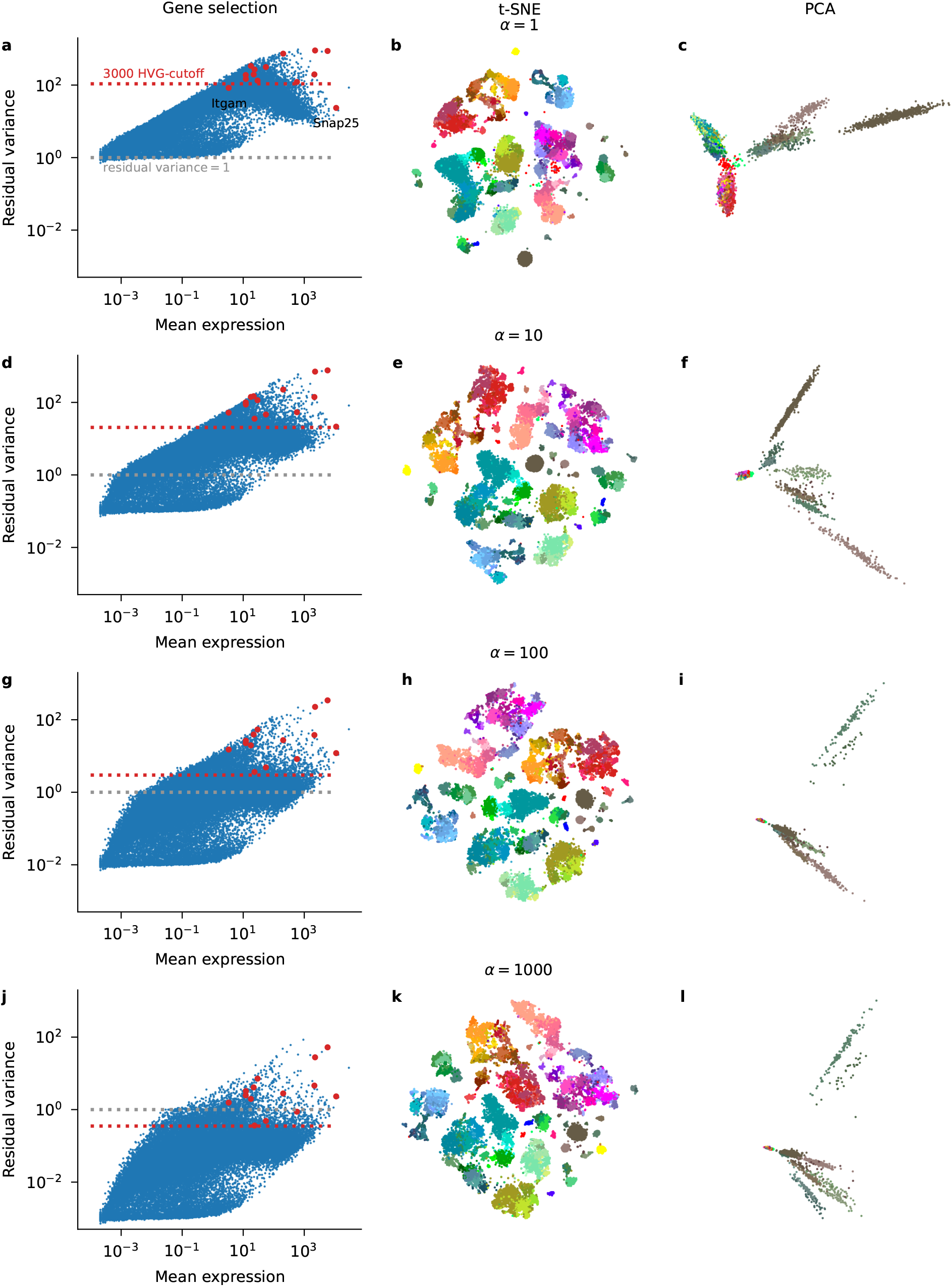
Influence of the amplification parameter *α*. Each row contains a reproduction of Figure 3a– b for various values of *α*_*Z*_. The first row corresponds to the NB model without amplification, used for UMI data (Lause et al., 2021). Gray line: indicates residual variance = 1, where most non-differentially expressed genes should lie if the model is correct. All t-SNEs used the same shared initialization (see Methods). Right column shows the first two principal components (PCs) of the compound Pearson residuals.

**Figure S7:**
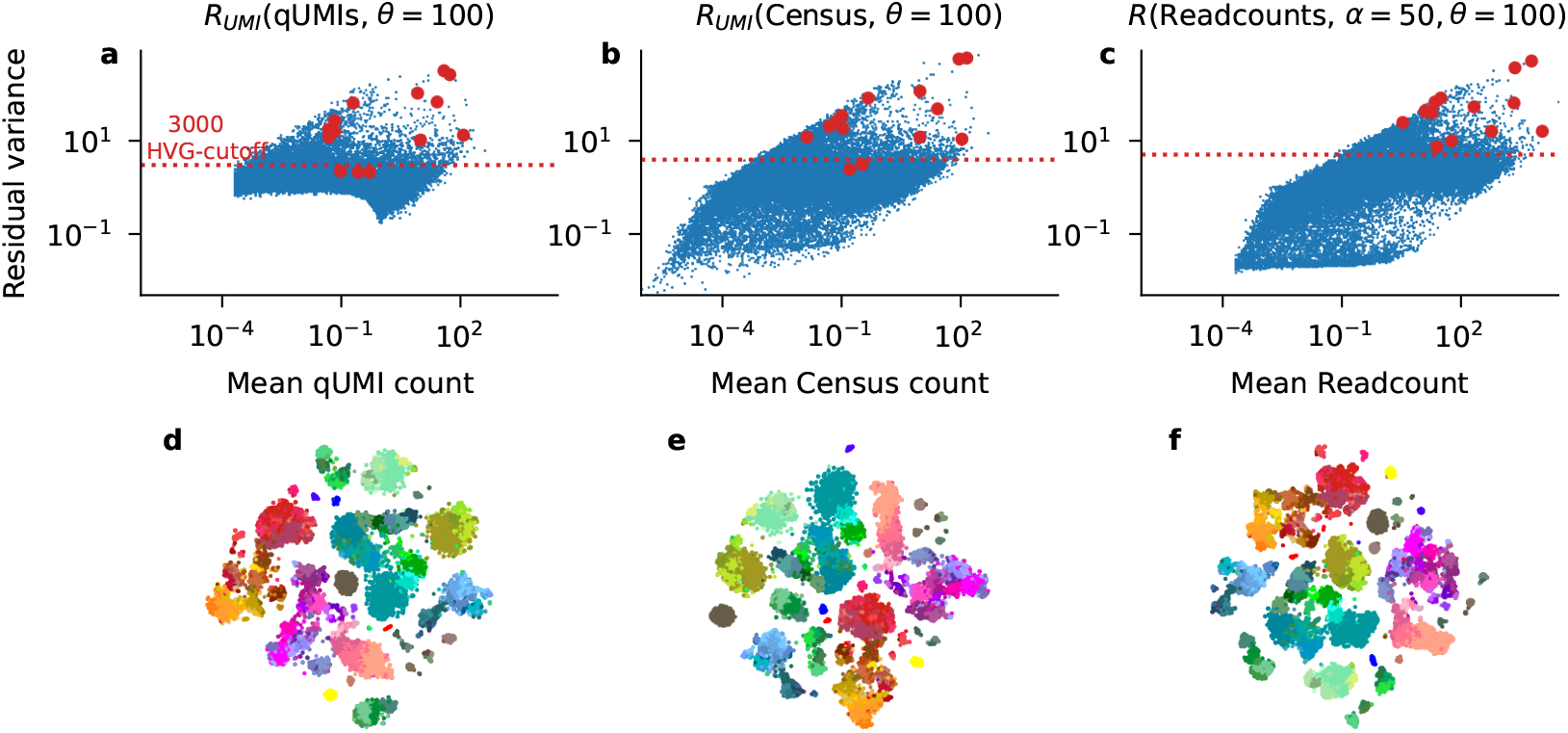
Comparing compound Pearson residuals to Census and qUMI. The same analysis as shown in Figure 3a–b for read counts from Tasic et al. (2018) processed with qUMIs (Townes and Irizarry, 2020) followed by UMI Pearson residuals (a, d); Census counts (Qiu et al., 2017) followed by UMI Pearson residuals (b, e); and compound Pearson residuals (our method) applied to the same set of genes (see Methods) (c, f).

**Figure S8:**
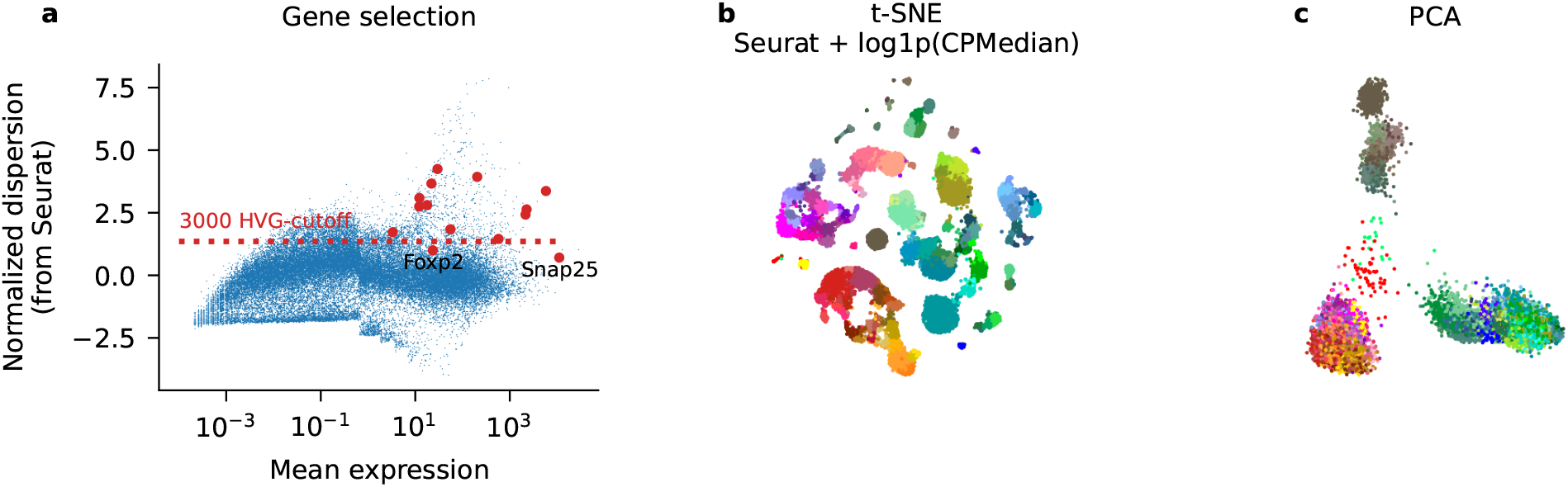
Pre-processing Smart-seq2 data with Scanpy default settings. The same dataset as in Figure S6, processed with scanpy 1.9.0 defaults for normalization (counts per median normalization with `normalize total()`, followed by `log1p()` transform) and gene selection (flavor=‘seurat’, Satija et al. (2015)) and PCA to 1 000 PCs. **a:** Seurat gene selection based on normalized dispersion. Note that two known markers (*Foxp2, Snap25*) are not among the top 3 000 genes selected by this method. The genes with highest normalized dispersion are markers of small, non-neural populations. **b:** t-SNE embedding of the pre-processed data. **c:** PCA embedding of the same, pre-processed data.

**Figure S9:**
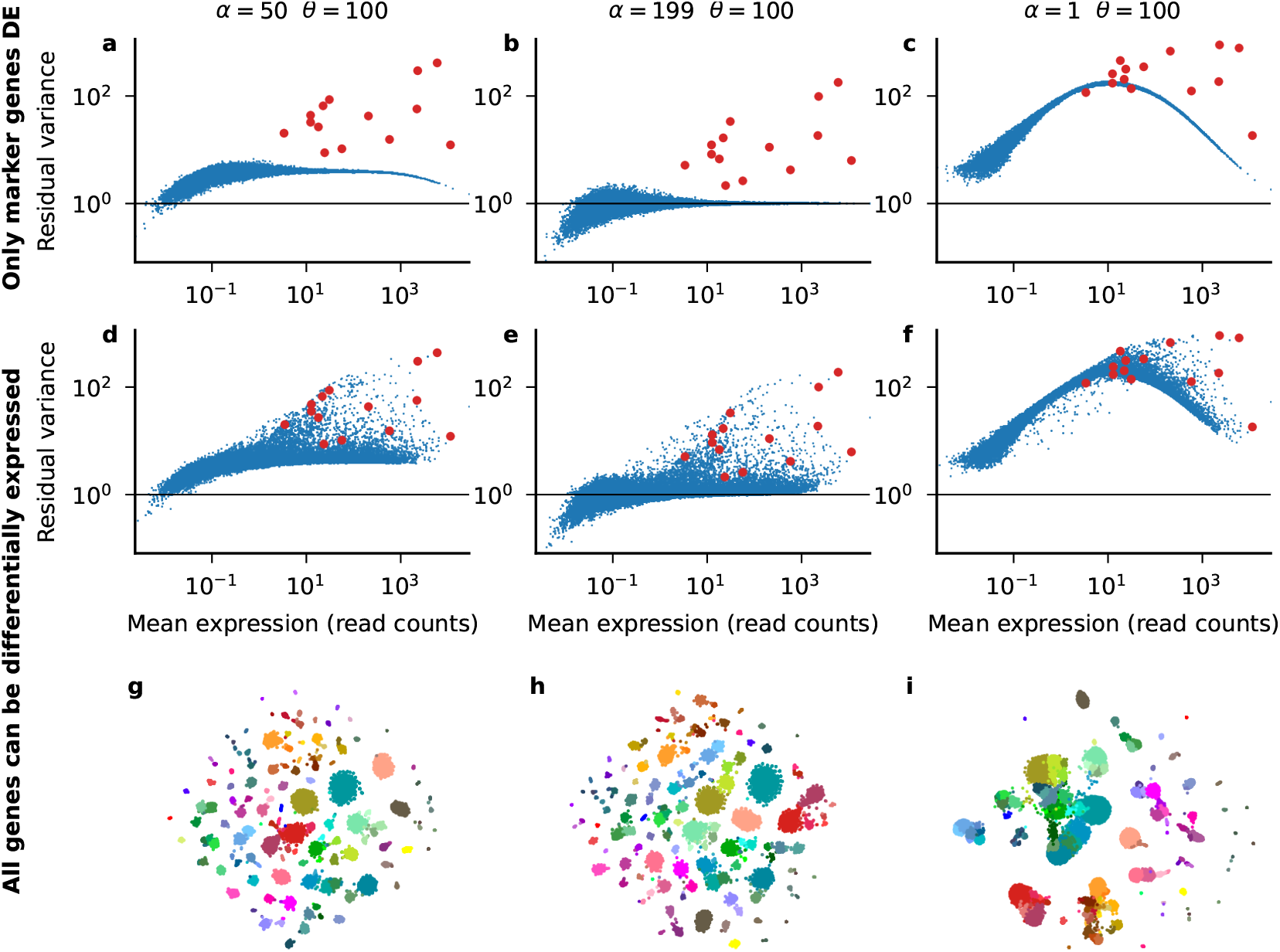
Compound Pearson residuals recover ground truth in realistic simulations. The same analysis as shown in Figure 3 for two simulated read count datasets that mirror the Tasic et al. (2018) cluster structure. Both simulated datasets were processed by compound Pearson residuals with three different settings: *α*_*Z*_ = 50 as in Figure 3 (left); *α*_*Z*_ = 199 which is the ground truth amplification factor used in this simulation (middle); *alpha*_*Z*_ = 1 corresponding to UMI Pearson residuals (right). **a–c:** Simulation I. Marker genes (red) were simulated with cluster-specific expression strengths from the Tasic et al. (2018) data, all other genes with their average expression strength across the whole dataset. Horizontal line indicates unit residual variance, expected for genes without differential expression. **d–i:** Simulation II. All genes were simulated with their cluster-specific expression strengths.

**Figure S10:**
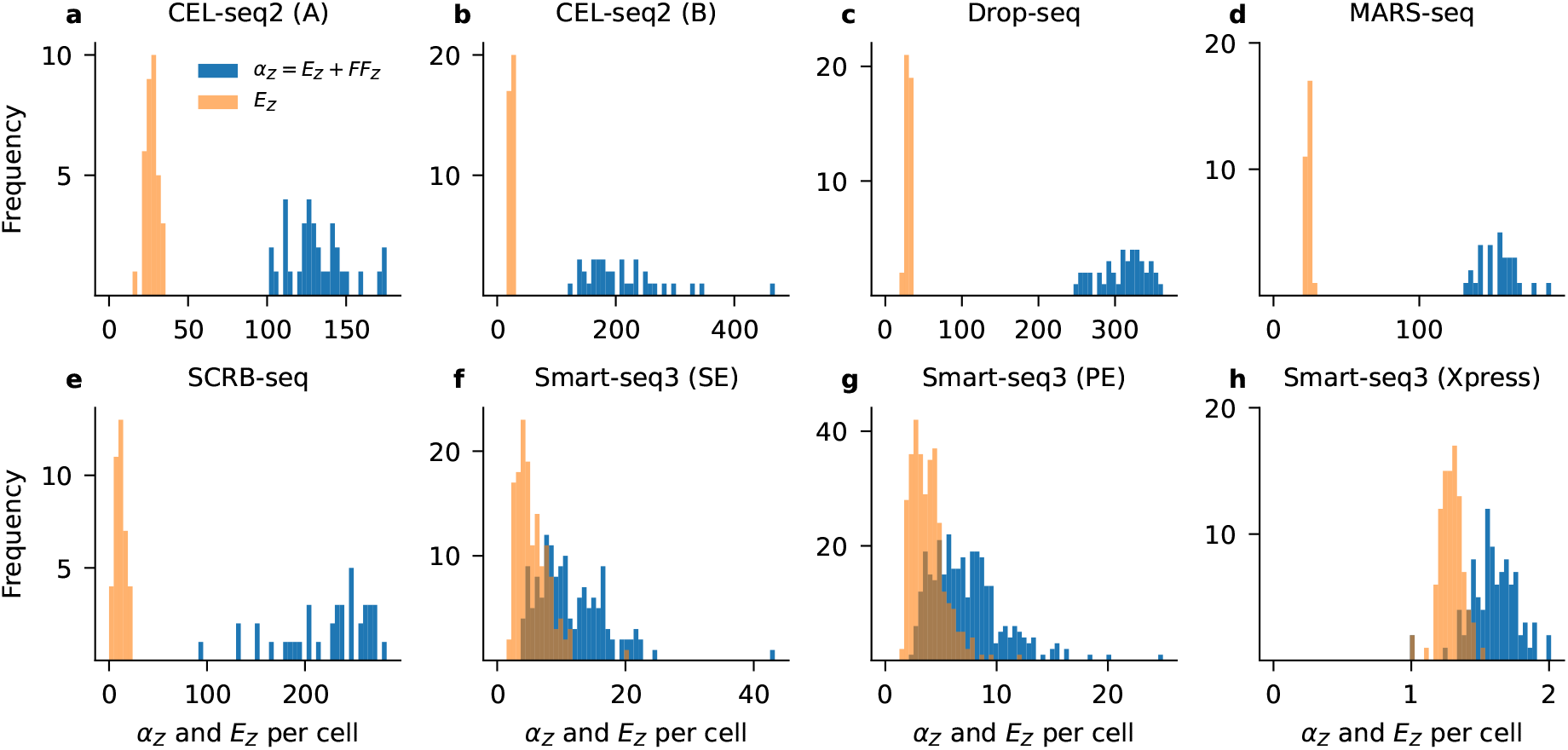
Cell-to-cell variability in *α*_*Z*_ and E[*Z*]. Each panel shows amplification statistics computed per cell for all sequencing platforms listed in Table 1. Only run A is shown unless otherwise indicated.

**Figure S11:**
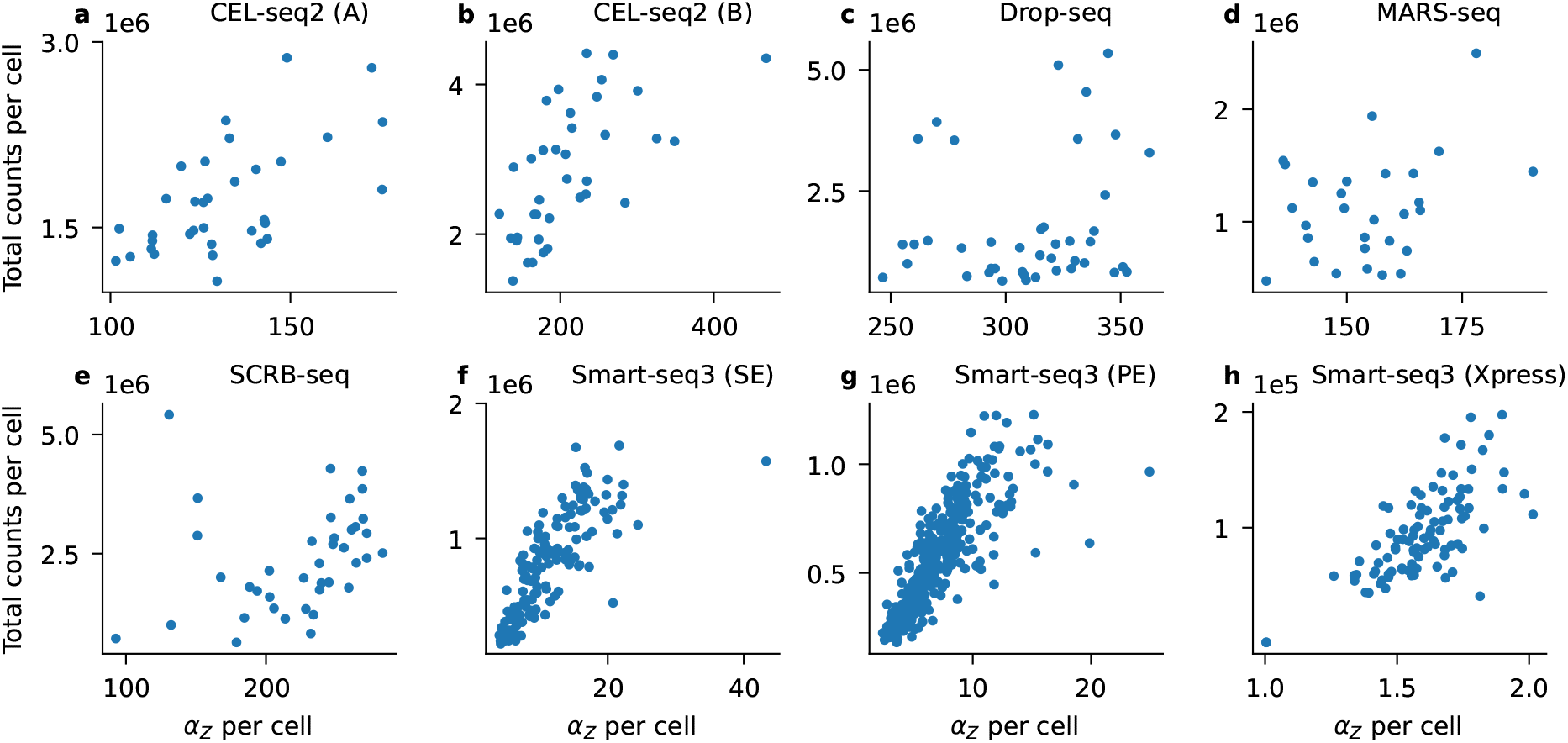
Total read counts are correlated with *α*_*Z*_. Each panel corresponds to one of the sequencing platforms listed in Table 1; each dot is a cell. Only run A is shown unless otherwise indicated.

